# RyR2 regulates store-operated Ca^2+^ entry, phospholipase C activity, and electrical excitability in the insulinoma cell line INS-1

**DOI:** 10.1101/2022.10.19.512717

**Authors:** Kyle E. Harvey, Shiqi Tang, Emily K. LaVigne, Evan P.S. Pratt, Gregory H. Hockerman

## Abstract

The ER Ca^2+^ channel ryanodine receptor 2 (RyR2) is required for maintenance of insulin content and glucose-stimulated insulin secretion, in part, via regulation of the protein IRBIT in the insulinoma cell line INS-1. Here, we examined store-operated and depolarization-dependent Ca^2+^entry using INS-1 cells in which either RyR2 or IRBIT were deleted. Store-operated Ca^2+^ entry (SOCE) stimulated with thapsigargin was reduced in RyR2^KO^ cells compared to controls, but was unchanged in IRBIT^KO^ cells. STIM1 protein levels were not different between the three cell lines. Basal and stimulated (500 μM carbachol) phospholipase C (PLC) activity was also reduced specifically in RyR2^KO^ cells. Insulin secretion stimulated by tolbutamide was reduced in RyR2^KO^ and IRBIT^KO^ cells compared to controls, but was potentiated by an EPAC-selective cAMP analog in all three cell lines. Cellular PIP_2_ levels were increased and cortical f-actin levels were reduced in RyR2^KO^ cells compared to controls. Whole-cell Ca_v_ channel current density was increased by 65% in RyR2^KO^ cells compared to controls, and barium current was reduced by acute activation of the lipid phosphatase pseudojanin preferentially in RyR2^KO^ cells over control INS-1 cells. Action potentials stimulated by 18 mM glucose were more frequent in RyR2^KO^ cells compared to controls, and insensitive to the SK channel inhibitor apamin. Taken together, these results suggest that RyR2 plays a critical role in regulating PLC activity and PIP_2_ levels via regulation of SOCE. RyR2 also regulates β-cell electrical activity by controlling Ca_v_ current density, via regulation of PIP_2_ levels, and SK channel activation.

## 1. Introduction

Temporal and spatial regulation of intracellular Ca^2+^ concentrations ([Ca^2+^]in) is critical for pancreatic β-cell function and survival [1], and aberrant Ca^2+^ handling is associated with type 2 diabetes [2]. Elevation of [Ca^2+^]_in_ in β-cells involves both influx of Ca^2+^ into cells via plasma membrane resident channels [1], and release of Ca^2+^ from the endoplasmic reticulum (ER) [3]. Influx of Ca^2+^ through voltage-gated Ca^2+^ (Ca_v_) channels is an essential step for maximal insulin secretion by pancreatic β-cells [4]. Ca^2+^ influx is amplified by activation of channels that release Ca^2+^ from internal stores through the process of Ca^2+^-induced Ca^2+^ release (CICR) [3, 5]. Voltage-gated Ca^2+^ entry is mediated through Ca_v_1.2, Ca_v_1.3, and Ca_v_2.1 in β-cells, and these ion channels are essential to insulin secretion and beta-cell survival [1, 4, 6]. While CICR is not essential for depolarization-induced insulin secretion, it is thought to amplify the effect [5, 7].

The two major classes of Ca^2+^ release channels residing on the ER membrane of β-cells are ryanodine receptors (RyR) and inositol 1,4,5-trisphosphate receptors (IP_3_R) [8]. RyR2 is expressed in human [9] and mouse β-cells [10], and is the predominant, if not sole, RyR functionally expressed in the rat insulinoma cell line INS-1 [11]. IP_3_R3 is the predominant IP_3_R expressed in human β-cells [8] and rat islets [12], but IP_3_R1 is predominant in mouse islets [13]. Dysregulation of Ca^2+^ homeostasis leads to ER stress and pancreatic β-cell death [14, 15]. ER Ca^2+^ depletion occurs under prolonged exposure to cytokines, fatty acids, and inhibition of sarcoplasmic/endoplasmic reticulum Ca^2+^ ATPase (SERCA)[16]. Store-operated Ca^2+^ entry (SOCE) is a mechanism by which ER Ca^2+^ stores are replenished by capacitative calcium entry, in contrast to depolarization-dependent Ca^2+^ influx [17]. Ca^2+^ release-activated channels (CRAC) mediate SOCE and are activated when ER Ca^2+^ is depleted. Upon a drop in ER Ca^2+^, the Ca^2+^ sensor STIM,1 present on the on the cytosolic face of the ER, translocates to ER-plasma membrane junctions. Upon translocation, STIM1 interact with Orai and TRPC1 allowing for influx of extracellular Ca^2+^ [18, 19]. SOCE is an essential process for refilling intracellular Ca^2+^ stores in non-excitable cells (i.e. lacking voltage-gated Ca^2+^ channels), but is also functionally significant in excitable cells [20]. Islets from type 2 diabetics were shown to have reductions in STIM1, and deletion of STIM1 from INS-1 pancreatic β-cells was shown to impair SOCE [21]. Pharmacological inhibition of SOCE inhibits glucose-stimulated insulin secretion, but inhibitors of SOCE have many off-target effects [21-23]. RyRs and IP_3_Rs may play a role in activation of SOCE. In pulmonary artery smooth muscle cells, stimulation of RyR2 can activate SOCE via a mechanism dependent on ER/SR Ca^2+^ depletion and a specific conformation of RyR2 [24]. IP_3_R may also play a role in the activation of SOCE in β-cells. Depletion of ER calcium stores by muscarinic receptor activation was sufficient to activate SOCE in INS-1E cells, and IP_3_R has direct and indirect interactions with various TRPC channels, including TRPC1 [22].

One potential role for SOCE, besides re-filling ER Ca^2+^ stores upon Ca^2+^ release via RyR and IP_3_R, is to provide Ca^2+^ to maintain phospholipase C (PLC) activity. PLCs are Ca^2+^-dependent enzymes that catalyze the hydrolysis of phosphatidyl inositol-4,5-bisphophate (PIP_2_) to inositol 1,4,5-trisphosphate (IP_3_) and diacylglycerol in the basal state, or upon stimulation by G_q_-coupled receptors [25]. SOCE plays a stimulatory role on prolonged PLC activity in response to the muscarinic receptor agonist carbachol in both MIN6 insulinoma cells and primary mouse pancreatic β-cells [26]. In these studies, 2-aminoethyl diphenylborate (2-APB), a small-molecule inhibitor of SOCE, acutely inhibited the late, sustained phase of PLC activity, but inhibition of membrane depolarization with diazoxide or inhibition of L-type VGCC with nifedipine was without effect. Another potential consequence of reduced PLC activity, particularly basal activity, is chronically increased PIP_2_ levels in the plasma membrane. An accumulation of PIP_2_ could have multiple effects on electrical activity in excitable cells since it has direct modulatory effects on ion channel activity [27], including that of Ca_v_ channel in β-cells [28].

We have previously demonstrated that crosstalk between RyR2 and IP_3_R is partially mediated through regulation of expression of IRBIT (IP_3_ <u>R</u>eceptor <u>B</u>inding protein released with Inositol 1,4,5 <u>T</u>risphosphate (a.k.a. AHCYL1)), as deletion of RyR2 from INS-1 cells leads to reduced levels of the protein IRBIT and subsequent dysregulation of IP_3_R [11]. Deletion of RyR2 also leads to marked reduction in insulin transcript, content, and glucose-stimulated secretion [11]. In the present study, we investigated the functional consequences of deletion of RyR2 and IRBIT on Ca^2+^ signaling via examination of SOCE, PLC activity, and electrical activity in control INS-1 cells and INS-1 cells in which RyR2 or IRBIT have been deleted using CRISPR/Cas9 gene editing. Our results suggest that RyR2 plays a crucial role in regulation of SOCE and PLC activity independent of IRBIT, as well as in the regulation of electrical activity in β-cells.

## 2. Materials and methods

### 2.1 Chemicals and Reagents

Fura-2 AM was from Molecular Devices (San Jose, CA). Xestospongin C and rapamycin were from Cayman Chemical (Ann Arbor, MI). Bethanechol was from Alfa Aesar (Haverhill, MA). Apamin was from Tocris Bioscience (Bristol, UK). All other reagents, unless otherwise indicated, were from Sigma-Aldrich (St. Louis, MO). Phalloidin conjugates were from Biotium (Fremont, CA). Plasmids encoding Pseudojanin (PJ) (Addgene plasmid # 37999), LYN11-FRB-CFP (Addgene plasmid # 38003), and PJ-DEAD (PJ-D) (Addgene plasmid # 38002) were gifts from Robin Irvine. GFP-C1-PLCdelta-PH was a gift of Tobias Meyer (Addgene plasmid # 21179).

### 2.2 Cell Lines

INS-1 cells [29] (gifted by Dr. Ming Li, Tulane University) were cultured in RPMI-1640 medium (Sigma-Aldrich, St. Louis, MO) supplemented with 10% fetal bovine serum (Qualified, Gibco), 11 mg/mL sodium pyruvate, 10 mM HEPES, 100 U/mL penicillin, 100 µg/mL streptomycin, and 50 µM mercaptoethanol at 37°C, 5% CO_2_. Construction and characterization of the RyR2^KO^ and IRBIT^KO^ cell lines were described previously [11].

### 2.3 Single-Cell Intracellular

*Ca*^*2+*^ *Assays*-INS-1, RyR2^KO^, and IRBIT^KO^ cells were plated in a poly-D-lysine coated 4-chambered 35 mm glass bottom tissue culture dish (Cellvis, Mountain View, CA). Cells were incubated overnight in RPMI-1640 medium at 37°C, 5% CO_2_. Cells were washed twice with PBS and loaded with 3 µM Fura-2 AM (Thermo Fisher, Waltham, MA) diluted in a modified Krebs-Ringer buffer solution [KRBH: 134 mM NaCl, 3.5 mM KCl, 1.2 mM KH_2_PO_4_, 0.5 mM MgSO_4_, 1.5 mM CaCl_2_, 5 mM NaHCO_3_, 10 mM HEPES (pH 7.4)] supplemented with 0.05% fatty acid free BSA at room temperature for 1 hour. The KRBH-containing Fura-2 AM was removed, cells were washed twice with KRBH, then equilibrated in KRBH alone or KRBH containing a 2X concentration of indicated inhibitors for 30 minutes at room temperature. The 4-chambered 35 mm dish was mounted on a chamber attached to the stage of an Olympus IX50 inverted microscope equipped with a PlanApo 20X objective lens (0.95 na). Cells were stimulated with the indicated stimulus at a 2X concentration. Cells were alternatively excited at 340/11 nm and 380/20 nm wavelengths using a bandpass filter shutter (Sutter Instruments, Novato, CA) and changes in intracellular Ca^2+^ were measured by recording the ratio of fluorescence intensities at 508/20 nm in time lapse with a time interval of 0.6 seconds using a Clara CCD camera (Andor Technology, Belfast, Ireland). Background subtraction from the raw 340/11 nm and 380/20 nm wavelengths was performed, then isolated single cells were selected and traced as regions of interest (ROIs) and the 340/11 nm/380/20 nm ratios were measured for each ROI using MetaMorph image analysis software (Molecular Devices, San Jose, CA). Single-cell Ca^2+^ traces were normalized to their respective baseline intracellular Ca^2+^ level obtained by averaging the 340/11 nm/380/20 nm ratios during the first minute of each experiment in the absence of stimulus. Ca^2+^ transients are plotted as normalized 340/11 nm/380/20 mn ratios against time.

### 2.4 Intracellular Ca^2+^ Measurements in 96-well Plates

INS-1, RyR2^KO^, and IRBIT^KO^ cells were plated at 70-90% confluency in black-walled 96-well tissue culture plates (Corning, Corning, NY) in RPMI-1640 medium and incubated overnight at 37°C, 5% CO_2_. Cells were washed twice with PBS and incubated with 100 µL 5 µM Fura-2 AM diluted in KRBH for 1 hour at room temperature. The KRBH containing Fura-2 AM was removed, cells were washed twice with KRBH, then equilibrated for 30 minutes at room temperature. To measure SOCE, baseline fluorescence was measured for 1 minute. Thapsigargin was then injected to a final concentration of 1 µM to deplete ER Ca^2+^ stores and fluorescence was measured for 20 minutes. 2-APB was co-injected with thapsigargin in some experiments to a final concentration of 100 µM to block SOCE. Finally, a final concentration of 2.5 mM CaCl_2_ was injected, and fluorescence measured for 10 minutes. To measure fluorescence, cells were alternatively excited at 340/11 nm and 380/20 nm (center/bandpass) and changes in intracellular Ca^2+^ concentrations were measured by recording the ratio of fluorescence intensities at 508/20 nm (15 second time interval) using a Synergy 4 multimode microplate reader (BioTek, Winooski, VT). Traces were normalized to their respective baseline intracellular Ca^2+^ level obtained by averaging the 340/11 nm/380/20 nm ratios of each experiment in the absence of stimulus. Ca^2+^ transients are plotted as normalized 340/11 nm/380/20 nm ratios against time.

### 2.5 IP_1_ HTRF Assays

INS-1, RyR2^KO^, and IRBIT^KO^ cells were plated at approximately 200,000 cells/well in a white opaque 96-well tissue culture plate (Corning, Corning, NY) and incubated overnight in low glucose RPMI-1640 medium at 37°C, 5% CO_2_. Cells were washed twice with PBS and incubated in a pre-stimulation buffer [10 mM HEPES, 1 mM CaCl_2_, 0.5 mM MgCl_2_, 4.2 mM KCl, 146 mM NaCl (pH 7.4)] for 1 hour at 37°C, 5% CO_2_. The pre-stimulation buffer was decanted, and stimulants and/or inhibitors at the indicated concentrations were added to the cells in the same pre-stimulation buffer supplemented with 50 mM LiCl to inhibit inositol monophosphate degradation and incubated for 1 hour at 37°C, 5% CO_2_. Accumulation of IP_1_ was measured using the IP-One Gq Homogenous Time-Resolved Fluorescence (HTRF) kit from Perkin Elmer (Waltham, MA), per the manufacturer’s instructions. The IP_1_ concentration of each sample was interpolated by comparison to a standard curve of known IP_1_ concentrations.

### 2.6 Insulin Secretion Assays

INS-1, RyR2^KO^, and IRBIT^KO^ cells were plated at 70–90% confluency in 96-well plates (Corning) in RPMI-1640 media and incubated overnight at 37 °C, 5% CO_2_. 16–24 h prior to assay, cells were incubated in serum-free, low glucose RPMI-1640 media supplemented with 0.1% fatty acid-free BSA overnight at 37 °C, 5% CO_2_. Cells were washed once with PBS and pre-incubated with 100 μL fatty acid-free KRBH alone or containing the working concentration of ESCA for 30 min at 37 °C, 5% CO_2_. After 30 min, KRBH was removed and replaced with either 100 μL KRBH or KRBH containing the indicated concentrations of stimulants, and cells were stimulated for 30 min at 37 °C, 5% CO_2_.

Supernatants were collected and stored at -20 °C until assayed. Cells were lysed in 50 μL ice-cold modified RIPA lysis buffer (100 mM NaCl, 25 mM Tris pH 8, 1% Triton X-100, 0.5% sodium deoxycholate, 0.1% SDS, 5 mM MgCl_2_, 1 mM CaCl_2_) supplemented with 10 μg/mL DNAse I and protease inhibitors (1 mM 4-(2-aminoethyl) benzenesulfonyl fluoride hydrochloride, 800 nM aprotinin, 50 μM bestatin, 15 μM E-64, 20 μM leupeptin, and 10 μM pepstatin A). Protein content of lysates was measured using the Pierce BCA Protein Assay Kit (Thermo Fisher) per the manufacturer’s instructions. Insulin measurements were performed with Insulin High-Range assay kits (Perkin Elmer).

### 2.7 Total Internal Reflection Fluorescence Microscopy (TIRFm)

INS-1 cells were transfected with 500 ng GFP-C1-PLCdelta-PH [30] in 12-well plates using LipoJet (Signagen, Frederick, MD) per the manufacturer’s protocol. 24 for hours after transfections, cells were split into poly-D-lysine coated 4-chambered 35 mm glass bottom dishes. Cells were imaged the following morning using a Nikon Ti2-E Inverted Microscope equipped with a TIRF illuminator and a Perfect Focus System (PFS) using a 100x Plan Apo Lambda oil objective (NA 1.4) (Nikon, Tokyo, Japan). Cells were preincubated for 30 min in KRBH containing 2.5 mM glucose at 37°C and 5% CO_2_. After 30 min, cells were imaged at room temperature. Images were acquired at a 1 second interval, 200 msec exposure, on a ORCA-FLASH4.0 digital CMOS camera (Hamamatsu, Hamamatsu, Japan). A 1-minute baseline was recorded prior to addition of 2x concentration of carbachol +/-2-APB, 100 µM final concentration for both compounds.

### 2.8 Ca_v_ Channel Current Density measurements

INS-1, RyR2^KO^, and IRBIT^KO^ cells were plated at low confluency in 35 mm tissue culture dishes (Corning) and incubated overnight in RPMI-1640 medium, at 37°C, 5% CO_2_. Whole-cell patch clamp recordings were performed using an Axopatch 200B amplifier (Molecular Devices, San Jose, CA). Data were sampled at 10 kHz and filtered at 1 kHz (six-pole Bessel filter, -3dB). Patch pipettes were pulled from borosilicate glass capillaries (VWR, West Chester, PA), with a P-87 micropipette puller (Sutter instruments, Novato, CA). Pipettes were polished with an MF-830 microforge (Narishige, Tokyo, Japan) to an inside diameter of 3-5 µm. The extracellular solution contained (mM): 140 NaCl, 20 CsCl_2_, 10 BaCl_2_, 10 dextrose, 10 sucrose, 1 MgCl_2_, 10 HEPES, pH 7.4, ∼350 mOsm. The intracellular solution contained: 180 N-methyl-D-glucamine (NMDG), 12 phosphocreatine, 5 BAPTA, 4 MgCl_2_, 2 Na_2_ATP, 0.5 Na_3_GTP, 0.1 leupeptin, 40 HEPES, pH 7.3, ∼320 mOsm. Current traces were elicited with 100 msec steps to voltages ranging from -70 mV to +50 mV in 10 mV increments every two seconds, from a holding potential of -80 mV, with on-line P/-4 leak subtraction, using pClamp 10.7-11.2 software (Molecular Devices). Peak current (in picoAmperes (pA)) for each cell was divided by whole-cell capacitance (in picoFarads (pF)) to calculate current density (pA/pF). V_1/2_ activation values were determined by plotting normalized tail-current amplitudes versus the corresponding 100-millisecond depolarizing voltage steps from -70 to +50 mV, in 10 mV-increments, from a holding potential of -80 mV. The data were fit to the equation, I = 1/(1 + exp((V_1/2_ -V)/k)), where k is a slope factor. For experiments measuring the fraction of Ba^2+^ current blocked by nifedipine in INS-1, RyR2^KO^, and IRBIT^KO^ cells, 5 µM nifedipine was applied to cells under voltage-clamp using an RSC 160 perfusion system (Biologic, Grenoble, France) while stepping from a holding potential of -80 mV to +10 mV for 100 ms, every 20 seconds. For experiments using rapamycin activation of Pseudojanin [31], INS-1, and RyR2^KO^ cells were transfected with Lyn 11-FRP-CFP and either mRFP-FKBP-Pseudojanin active (PJ) or mRFP-FKBP-Pseudojanin phosphatase dead (PJ-D) using Lipojet transfection reagent (SignaGen, Frederick, MD). Rapamycin (1 µM) inhibition of current in transfected cells (identified by epifluorescence) was performed as described above for nifedipine block of current.

### 2.9 Perforated Patch Current Clamp recordings

Electrophysiological measurements of action potential frequencies were performed as described previously [5].

### 2.10 Data Analysis

Ca^2+^ imaging data, IP_1_ HTRF assay data, and insulin HTRF data were analyzed with Prism (9.3) software (GraphPad, San Diego, CA). Electrophysiological data were analyzed using Clampfit (10.7) software (Molecular Devices) and SigmaPlot (11.0) software (Systat Software, Palo Alto, CA). For statistical analysis, P < 0.05 was considered significant.

## 3. Results

### 3.1 RyR2 contributes to a rapid rise in [Ca^2^]_in_ upon membrane depolarization and suppresses IP_3_ receptor activation

Depolarizing stimuli have been shown to engage CICR in pancreatic β-cells, yet it is not clear whether release of Ca^2+^ from the ER is being mediated by RyR2 or IP_3_R [5]. Membrane depolarization of INS-1 cells with the K_ATP_ channel blocker tolbutamide (200 μM) results in a rapid peak followed by a sustained plateau in [Ca^2+^]_in_ as measured using Fura2-AM in a population of cells with a fluorescence-detecting plate reader, over two minutes (**Fig. 1A**). Pretreatment of INS-1 cells with 100μM ryanodine preferentially inhibited the fast peak of [Ca^2+^]_in_ stimulated by tolbutamide, largely leaving the sustained plateau in [Ca^2+^]_in_ intact (**Fig.1A**). Deletion of RyR2 from INS-1 cells using CRISPR/Cas9 gene editing abolished the fast peak in [Ca^2+^]_in_ under the same stimulus (**Fig.1B**). Measurement of changes in [Ca^2+^]_in_ upon stimulation with tolbutamide in single cells over 5 minutes revealed a delay in the time to peak [Ca^2+^]_in_ in RyR2^KO^ cells compared to control INS-1 (**Fig.1C,D)**. Since we’d previously found that RyR2 deletion results in down-regulation of the protein IRBIT [11], we examine the [Ca^2+^]_in_ response to tolbutamide in INS-1 cells in which IRBIT had been deleted using CRISPR/Cas9 gene editing (IRBIT^KO^ cells). We found that the time to peak [Ca^2+^]_in_ after stimulation with tolbutamide in IRBIT^KO^ cells was longer than in control INS-1 cells, but significantly shorter than in RyR2^KO^ cells. Area under the curve analysis of the tolbutamide-stimulated Ca^2+^ transients showed that deletion of RyR2 didn’t significantly alter the tolbutamide-stimulated Ca^2+^ integral, but deletion of IRBIT reduced the Ca^2+^ integral compared to RyR2^KO^ cells, but not control INS-1 cells. The pretreatment of cells with the IP_3_R blocker xestospongin c (1 μM) reduced the tolbutamide stimulated Ca^2+^ integral in RyR2^KO^ and IRBIT^KO^ cells, but not in control cells (**Fig.1E**). Tolbutamide is a member of the sulfonylurea class of antidiabetic drugs that is used clinically to stimulate insulin secretion, so we examined tolbutamide-stimulated insulin secretion in control INS-1, RyR2^KO^, and IRBITK^KO^ cells. Basal insulin secretion (2.5 mM glucose) was reduced in the RyR2^KO^ cells compared to control INS-1 cells, and insulin secretion stimulated by 200 μM tolbutamide was significantly reduced in both RyR2^KO^ and IRBIT^KO^ cells compared to control INS-1 cells (**Fig. 1F**). Activation of the cAMP effector protein EPAC2 is thought to potentiate secretion of insulin partially through its ability to mobilize intracellular Ca^2+^ [32]. Therefore, we examined the ability of the cell-permeable <u>E</u>PAC <u>s</u>elective <u>c</u>AMP <u>a</u>nalog CPT-2’-O-Me-cAMP-AM (ESCA; 5 μM) to potentiate tolbutamide-stimulated secretion. We found that ESCA significantly potentiated tolbutamide-stimulated insulin secretion in all three cell lines, but that the magnitude of insulin secretion stimulated by tolbutamide + ESCA was lower in RyR2^KO^ and IRBIT^KO^ cells compared to control INS-1 cells (**Fig. 1F**). Thus, deletion of RyR2 or IRBIT delays the peak of [Ca^2+^]_in_ stimulated by tolbutamide, and permits activation of IP_3_ receptors upon membrane depolarization in INS-1 cells. Further, deletion of RyR2 or IRBIT reduces insulin secretion stimulated by tolbutamide alone or by tolbutamide + ESCA, while deletion of RyR2 sharply reduces basal insulin secretion.

**Figure 1.**
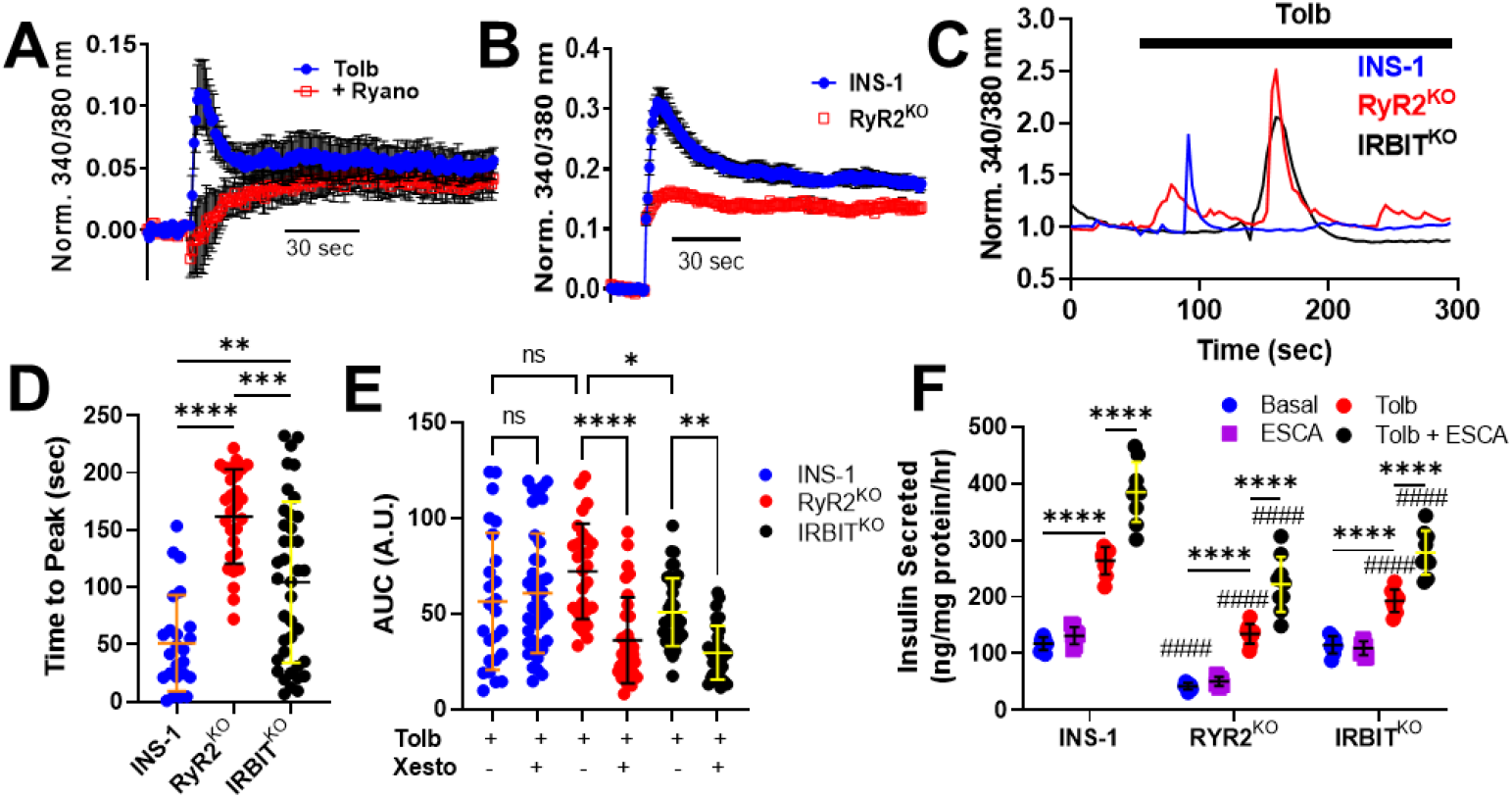
Deletion of RYR2 reshapes the [Ca^2+^]_in_ transient and impairs insulin secretion in response to tolbutamide-. **A)** Tolbutamide (200 µM) stimulates a biphasic rise in [Ca^2+^]_in_ in INS-1 cells. Preincubation with ryanodine (100 µM) selectively inhibits the initial peak in the [Ca^2+^]_in_ response. **B)** RyR2^KO^ cells stimulated with tolbutamide show selective loss of the rapid peak of [Ca^2+^]_in_ compared to control INS-1 cells. Experiments in A and B were performed in a 96-well format, over two minutes. Tolbutamide (200 µM) was injected after a 15 sec baseline was recorded. Each point is the mean of triplicate values shown ± SE. **C)** Example traces from single cell [Ca^2+^]_in_ imaging of a control INS-1 cell, an RyR2^KO^ cell, and an IRBIT^KO^ cell over 5 minutes. Tolbutamide was applied at 60 seconds. **D)** Time to greatest peak analysis of INS-1, RyR2^KO^, and IRBIT^KO^ cells. The time to peak was longer for both RyR2^KO^ and IRBIT^KO^ cells compared to INS-1 cell (\*\**P* < 0.01; \*\*\*\**P* < 0.0001 One-way ANOVA with Tukey’s post-hoc test). The time to peak was longer in RyR2^KO^ compared to IRBIT^KO^ cells (\*\*\**P* < 0.001) One-way ANOVA with Tukey’s multiple comparisons test. INS-1n = 24; RyR2^KO^: n = 28; IRBIT^KO^: n = 39. **E)** Area under the curve analysis (AUC) of the [Ca^2+^]_in_ response to tolbutamide. The AUC was significantly reduced in IRBIT^KO^ cells compared to RyR2^KO^ cells (*, *P* < 0.05), and 1 µM xesto reduced AUC in RyR2^KO^ and IRBIT^KO^ cells only (****, *P* < 00001; **, *P* < 0.01) One-way ANOVA with Tukey’s multiple comparisons test. INS-1: n = 25; INS-1 + xesto: n = 40; RyR2^KO^: n = 28; RyR2^KO^ + xesto: n = 32; IRBIT^KO^: n = 37; IRBIT^KO^ + xesto: n = 30. **F)** Deletion of RyR2 or IRBIT reduces tolbutamide-stimulated insulin secretion compared to INS-1 cells, and reduces insulin secretion stimulated by tolbutamide + ESCA (^####^*P* < 0.0001). Basal insulin secretion is reduced in RyR2^KO^ cells compared to INS-1 cells (^####^, *P* < 0.0001). In all cases, tolbutamide stimulated an increase in insulin secretion over basal (2.5 M glucose; ****, *P* < 0.0001) and tolbutamide + ESCA stimulated an increase in insulin secretion over tolbutamide alone (****, *P* < 0.0001). n = 9 for all conditions. Two-way ANOVA with Tukey’s multiple comparisons test.

### 3.2 RyR2 regulates PLC activity in INS-1 cells

Parasympathetic innervation of the pancreas is thought to enhance insulin secretion after meals through activation of muscarinic receptors present on β-cells [33]. Muscarinic receptors M3 and M5 are found in β-cells, and are coupled to Gq signaling, upstream of PLC activation [34]. Therefore, we examined the ability of the muscarinic agonist bethanechol to elevate [Ca^2+^]_in_ in RyR2^KO^, IRBIT^KO^, and control INS-1 cells (**Fig. 2A**). We compared three concentrations of bethanechol, 1, 5, and 50 µM, to determine if the enhanced activation of IP_3_R observed during tolbutamide stimulation in RyR2^KO^ and IRBIT^KO^ cells enhances bethanechol stimulation of [Ca^2+^]_in_ increases. The Ca^2+^ integrals stimulated by 1 µM or 5 µM bethanechol didn’t differ among the three cell lines, and weren’t different from those observed in the presence of 50 µM bethanechol + 100 µM atropine (**Fig. 2B**). In contrast, 50 µM bethanechol, in the absence of atropine, stimulated a Ca^2+^ transient in all three cell lines that was significantly greater than that stimulated by 1 or 5 µM bethanechol, and the Ca^2+^ integral was greater in both RyR2^KO^ and IRBIT^KO^ cells than in control INS-1 cells (**Fig. 2B**). In addition, the Ca^2+^ integral stimulated by 50 µM bethanechol in IRBIT^KO^ cells was greater than that stimulated in RyR2^KO^ cells (**Fig. 2B**). Thus, deletion of IRBIT or RyR2 enhances the stimulation of [Ca^2+^]_in_ by 50 µM bethanechol in INS-1 cells, but deletion of IRBIT leads to a greater [Ca^2+^]_in_ response to 50 µM bethanechol than deletion of RyR2.

**Figure 2.**
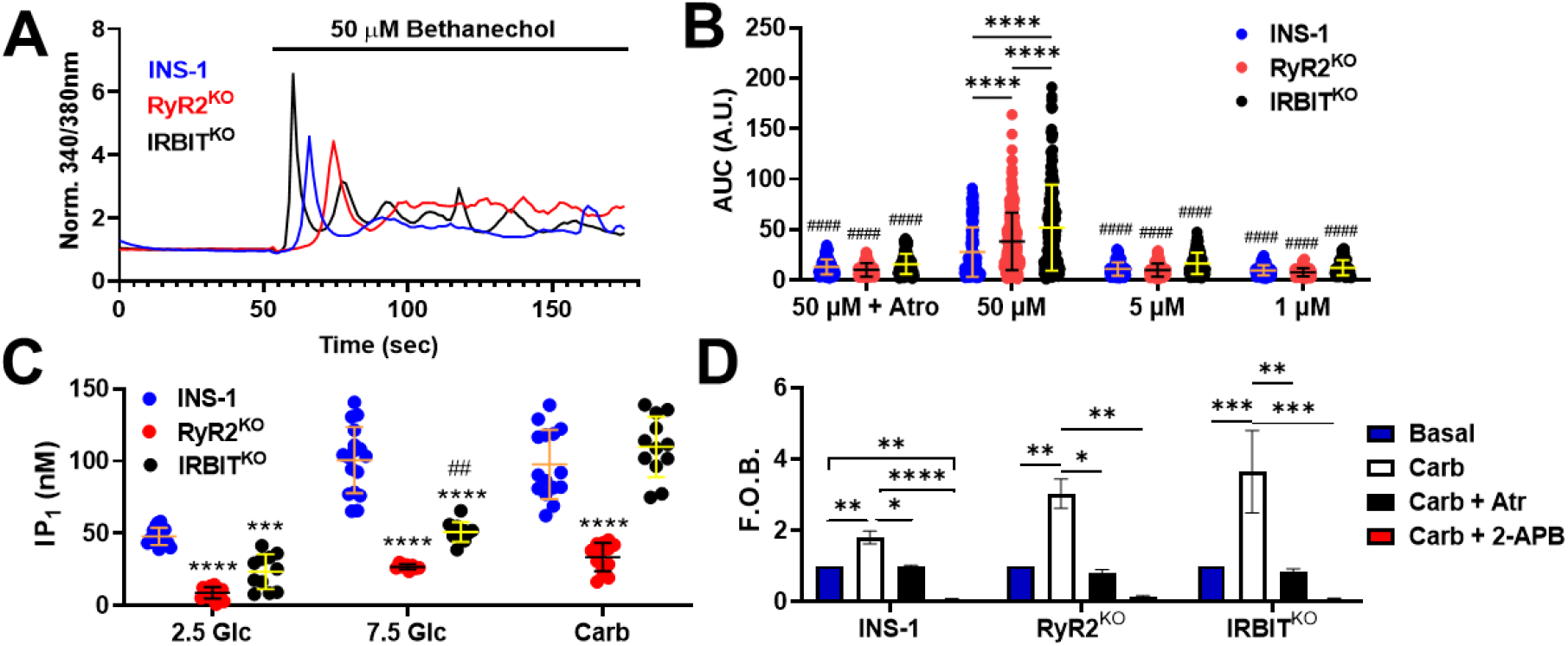
Deletion of RYR2 and IRBIT reduce stimulated and basal PLC activity-. **A)** Example traces from single-cell [Ca^2+^]_in_ imaging of a control INS-1 cell, an RyR2^KO^ cell, and an IRBIT^KO^ cell over 3 minutes. Bethanechol was applied at 55-60 seconds. **B)** Quantification of the increase in [Ca^2+^]_in_ stimulated by the muscarinic agonist bethanechol (AUC). 50 µM bethanechol stimulates a greater increase in [Ca^2+^]_in_ in IRBIT^KO^ and RyR2^KO^ cells than in INS-1cells (****, *P* < 0.0001), and a greater increase in IRBIT^KO^ cells than in RyR2^KO^ cells (****, *P* < 0.0001). Stimulation of [Ca^2+^]_in_ by 1 µM or 5 µM bethanechol is not different between the three cell lines, but in each case, is different from that stimulated by 50 µM bethanechol (^####^, *P* < 0.0001). Atropine (100 µM) significantly reduced the [Ca^2+^]_in_ response to 50 µM bethanechol in all cell lines (^####^, *P* < 0.0001) INS-1: 50 µM + Atro n = 175; 50 µM n = 143; 5 µM n = 153, 1 µM n = 132. Ry2R^KO^: 50 µM + Atro n = 164; 50 µM n = 293; 5 µM n = 159, 1 µM n = 111. IRBIT^KO^: 50 µM + Atro n = 106; 50 µM n = 195; 5 µM n = 126, 1 µM n = 102. Outlier analysis was performed using the ROUT method, with Q =1%. Outliers were removed before statistical analysis. **C)** IP_1_ assay of basal and stimulated PLC activity. Basal (2.5 M glucose) and 7.5 M glucose-stimulated PLC activity was reduced in RyR2^KO^ and IRBIT^KO^ cells compared to INS-1 cells (***, *P* < 0.001; ****, *P* < 0.0001). Glucose-stimulated IP_1_ accumulation was greater in IRBIT^KO^ cells than in RyR2^KO^ cells (^##^, *P* < 0;01). In contrast, carbachol (500 µM) stimulated IP_1_ accumulation was significantly reduced in RyR2^KO^ cells compared to both control cells and IRBIT^KO^ cells (****, *P* < 0.0001). Data are shown as mean ±SD. INS-1: 2.5 Glc n= 16, 7.5 Glc n = 17, Carb n = 15; RyR2^KO^: 2.5 Glc n = 13, 7.5 Glc n = 8, Carb n = 14; IRBIT^KO^: 2.5 Glc n = 11, 7.5 Glc n = 11, Carb n = 12. Data for INS-1 cells and RyR2^KO^ cells were previously published in [11], and are shown for comparison with data from IRBIT^KO^ cells. **D)** IP_1_ accumulation stimulated by carbachol in INS-1, RyR2^KO^, and IRBIT^KO^ cells is inhibited by 100 µM atropine and 100 µM 2-APB (*, *P* < 0.05; **, *P* < 0.01; ***, *P* < 0.001; ****, *P* < 0.0001). Data are shown as mean ± SD. N = 4 for each condition in all cell lines. Two-way ANOVA with Tukey’s multiple comparisons test.

We next examined the ability of 7.5 mM glucose and the muscarinic agonist carbachol (500 µM) to stimulate phospholipase C activity in INS-1, RyR2^KO^, and IRBIT^KO^ cells. For these experiments, we used a homogeneous time-resolved FRET assay for inositol-1-phosphate (IP_1_). In the presence of lithium chloride, IP_1_ is a stable metabolite of IP_3_ and can serve as a surrogate marker for PLC activation and IP_3_ generation [35]. IP_1_ accumulation in 2.5 mM (basal) and 7.5 mM glucose were reduced in both RyR2^KO^ and IRBIT^KO^ cells compared to control INS-1 cells, but IP_1_ accumulation was significantly greater in IRBIT^KO^ cells than in RyR2^KO^ cells in the presence of 7.5 mM glucose (**Fig. 2C**). In contrast, IP_1_ accumulation stimulated by carbachol was only reduced in RyR2^KO^ cells compared to control INS-1 cells (**Fig. 2C**). To ensure that the PLC activity stimulated by carbachol was, in fact, mediated by muscarinic receptor activation, performed a set of experiments that included inhibition of the response with 100 µM atropine. We found that atropine reduced the accumulation of IP_1_ stimulated by carbachol to a level not different from basal levels (2.5 mM glucose) in all three cell lines (**Fig. 2D**). Store-operated calcium entry (SOCE) is proposed to provide Ca^2+^ to support PLC activity [26]. Accordingly, the application of the SOCE inhibitor 2-APB (100 μM) inhibited carbachol-stimulated IP_1_ accumulation below basal levels in all three cell lines (**Fig. 2D**).

Since muscarinic receptor-stimulated PLC activity appeared to be reduced in RyR2^KO^ cells, but not IRBIT^KO^ cells, as assessed with the IP_1_ assay, we examined the PIP_2_ levels in control INS-1, RyR2^KO^, and IRBIT^KO^ cells. Using immunofluorescence (primary antibody against PIP_2_) of fixed cells, we found that cellular PIP2 levels were significantly elevated in both IRBIT^KO^ and RYR2^KO^ cells compared to control INS-1 cells (**Fig. 3A**). Therefore, lack of substrate (i.e. PIP_2_) doesn’t appear to account for the decreased muscarinic receptor-stimulated PLC activity in RyR2^KO^ cells. To complement the results of the IP_1_ assay, we measured PLC activity in live cells using a probe consisting of the Pleckstrin homology domain of PLC delta fused to GFP (GFP-C1-PLCdelta-PH) [30]. This probe binds to PIP_2_, and can be imaged selectively in the plasma membrane using total internal reflection fluorescence microscopy (TIRFm). Acute stimulation of PLC activity using 100 µM carbachol resulted in a rapid decrease in GFP fluorescence intensity detected by TIRFm in INS-1 cells expressing GFP-C1-PLCdelta-PH, reflective of the acute decrease in PIP_2_ concentrations in the plasma membrane as it’s converted to diacylglycerol and IP_3_ (**Fig. 3C, D**). Pre-treatment of INS-1 cells expressing GFP-C1-PLCdelta-PH with 100 µM 2-APB strongly inhibited the decrease in plasma membrane GFP fluorescence intensity stimulated by carbachol (**Fig. 3E**). In contrast, carbachol stimulated a significantly smaller decrease in plasma membrane GFP fluorescence intensity in RyR2^KO^ cells expressing GFP-C1-PLCdelta-PH compared to that detected in control INS-1 cells (**Fig. 3D, E**). Further, the decrease in GFP-C1-PLCdelta-PH plasma membrane fluorescence intensity stimulated by carbachol in RyR2^KO^ cells was not significantly inhibited by 100 µM 2-APB (**Fig. 3E**). Thus, cellular PIP_2_ levels are increased in RyR2^KO^ and IRBIT^KO^ cells compared to control INS-1 cells, but PLC activity stimulated by carbachol is reduced in RyR2^KO^ cells compared to control INS-1 cells, and is insensitive to 2-APB.

**Figure 3.**
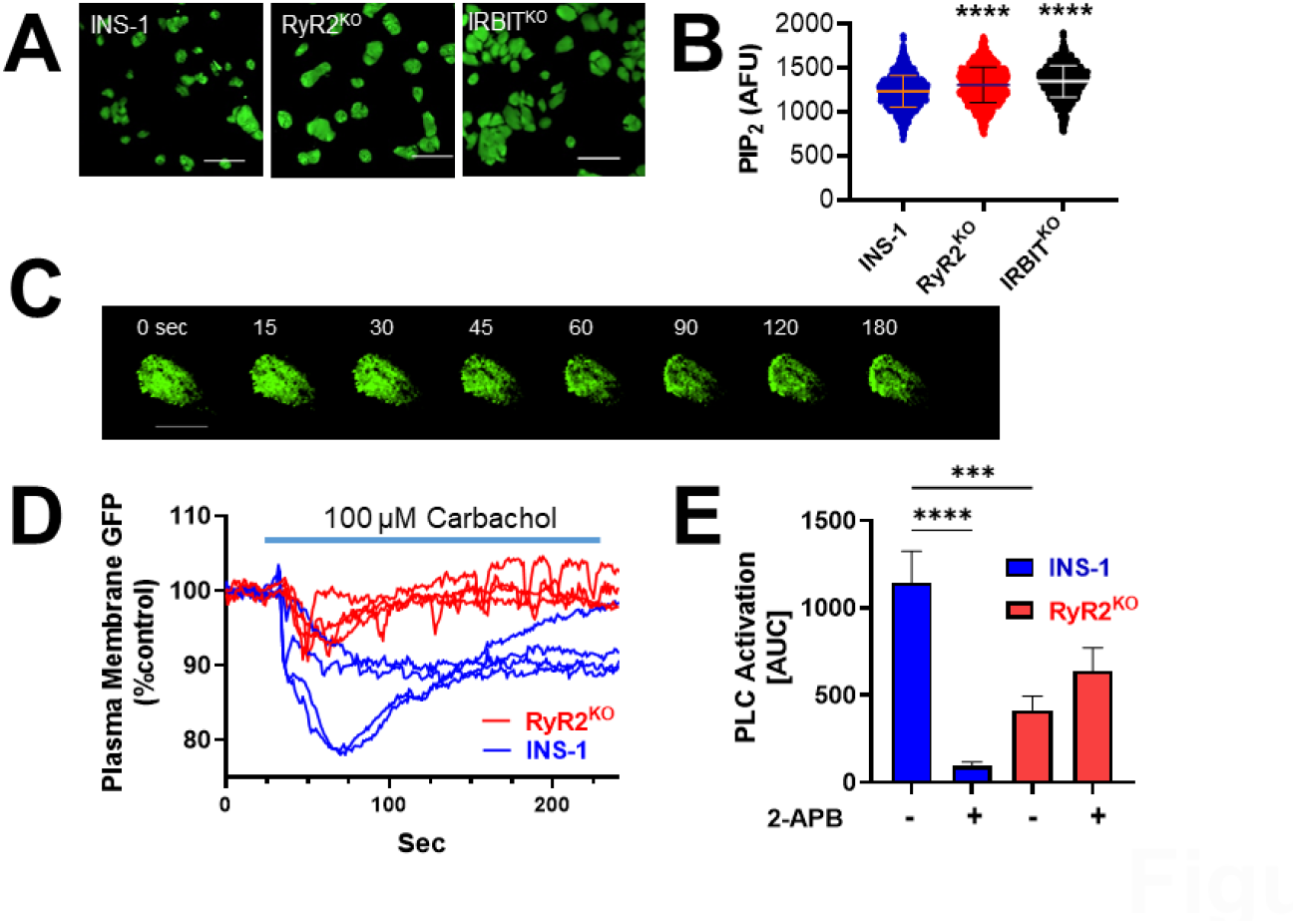
Modulation of PIP_2_ levels by PLC activity in INS-1 and RyR2^KO^ cells-. **A)** Micrographs of fixed INS-1, RyR2^KO^, and IRBIT^KO^ cells stained with antibodies to PIP_2_ and IgG-k binding protein conjugated to CFL 488. Scale bars = 50 µ. **B)** Quantification of fluorescence intensity of PIP_2_ immunostaining in control INS-1 cells, RyR2^KO^ cells, and IRBIT^KO^ cells. PIP_2_ staining was greater in RyR2^KO^ cells and IRBIT^KO^ cells compared to control INS-1 cells (****, *P* < 0.0001). INS-1 cells: n = 1294; RyR2^KO^ cells: n = 1377; IRBIT^KO^ cells n = 1650. Data are shown as mean ±SD. One-way ANOVA with Dunnett’s multiple comparisons test. Data for control INS-1 cells and RyR2^KO^ cells were previously published in [11] and are shown for comparison with data from IRBIT^KO^ cells. **C)** Tie lapsed TIRF images of GFP-C1-PLCdelta-PH localization to the plasma membrane upon stimulation with 100 µM carbachol in a control INS-1 cell. Scale bar = 10 µ. **D)** Example traces showing decreases in plasma membrane GFP-C1-PLCdelta-PH fluorescence intensity in response to carbachol in living INS-1 and RyR2^KO^ cells. **E)** Quantification (AUC) of the decrease in plasma membrane GFP-C1-PLCdelta-PH fluorescence in control INS-1 cells and RyR2^KO^ cells in the presence or absence of 100 µM 2-APB. Carbachol stimulated a greater decrease in PH-PLCδ-GFP fluorescence intensity in control INS-1 cells than in RyR2^KO^ cells (***, *P* < 0.001). 2-APB significantly inhibited this decrease in control INS-1 cells (****, *P* < 0.0001), but not in RyR2^KO^ cells. INS-1 cells + Carb: n = 27; RyR2^KO^ cells + Carb: n = 25; INS-1 cells + Carb + 2-APB: n = 18; RyR2^KO^ + Carb + 2-APB: n = 13. One-way ANOVA with Tukey’s multiples comparisons test.

### 3.3 SOCE is impaired by deletion of RyR2, but not IRBIT

SOCE is an important mechanism that cells utilize to replenish internal stores of calcium, but is also implicated in maintenance of PLC activity during stimulation [26]. Therefore, we examined SOCE in RyR2^KO^, IRBIT^KO^, and control INS-1 cells to determine if a deficit in SOCE might explain the sharply decreased PLC activity in RyR2^KO^ cells. In this assay, fura2-AM was used to measure changes in [Ca^2+^]_in_ as cells were depleted of the ER Ca^2+^ by injection of 1 μM thapsigargin in the absence of extracellular calcium to activate the SOCE pathway, and twenty minutes later, 2.5 mM extracellular Ca^2+^ re-introduced to initiate Ca^2+^ influx via SOCE (**Fig. 1A**). The magnitude of the response was quantified by integrating the [Ca^2+^]_in_ over time for the 10 minutes after addition of 2.5 mM Ca^2+^ **(Fig. 4D**). In INS-1 cells, this protocol produced a robust SOCE response that was strongly inhibited by pretreatment of cells with 100 µM 2-APB (**Fig. 1A, D**). In contrast, the magnitude of the SOCE response detected with this protocol was significantly reduced in RyR2^KO^ cells, and was insensitive to 2-APB (**Fig. 4B, D**). However, the magnitude of the SOCE response in IRBIT^KO^ cells was not different from control INS-1 cells, and was strongly inhibited 2-APB (**Fig. 4C, D**). STIM1 is an essential component of SOCE, and decreases in STIM1 lead to impairments in SOCE [20, 21]. Impairments in SOCE and reduced expression of STIM1 have been associated with β-cell dysfunction [21]. Therefore, we performed semi-quantitative immunoblotting to assess protein levels of STIM1 in RyR2^KO^, IRBIT^KO^, and control INS-1 cells, and found no difference between the three cell lines (**Fig 4 E, F**). 2-APB is reported to have multiple targets [36], so we performed the SOCE assays under conditions that minimized exposure of the cells to 2-APB. In this modified assay, 2-APB was co-applied to cells with thapsigargin, and extracellular Ca^2+^ was added back to cells 5 minutes after thapsigargin + 2-APB application. The resulting [Ca^2+^]_in_ responses in the presence or absence of 2-APB were quantified as before. This modified assay essentially reproduced the results in **Fig. 4** (i.e. RyR2^KO^ cells have impaired SOCE compared to INS-1 cells and IRBIT^KO^ cells) except that SOCE in RyR2^KO^ cells was inhibited by 2-APB (**Fig. S1**). Thus, deletion of RyR2 leads to reduced SOCE in INS-1 cells, while deletion of IRBIT is without effect on SOCE. This deficit in SOCE in RyR2^KO^ cells is likely not due to a reduction of STIM1 protein levels.

**Figure 4.**
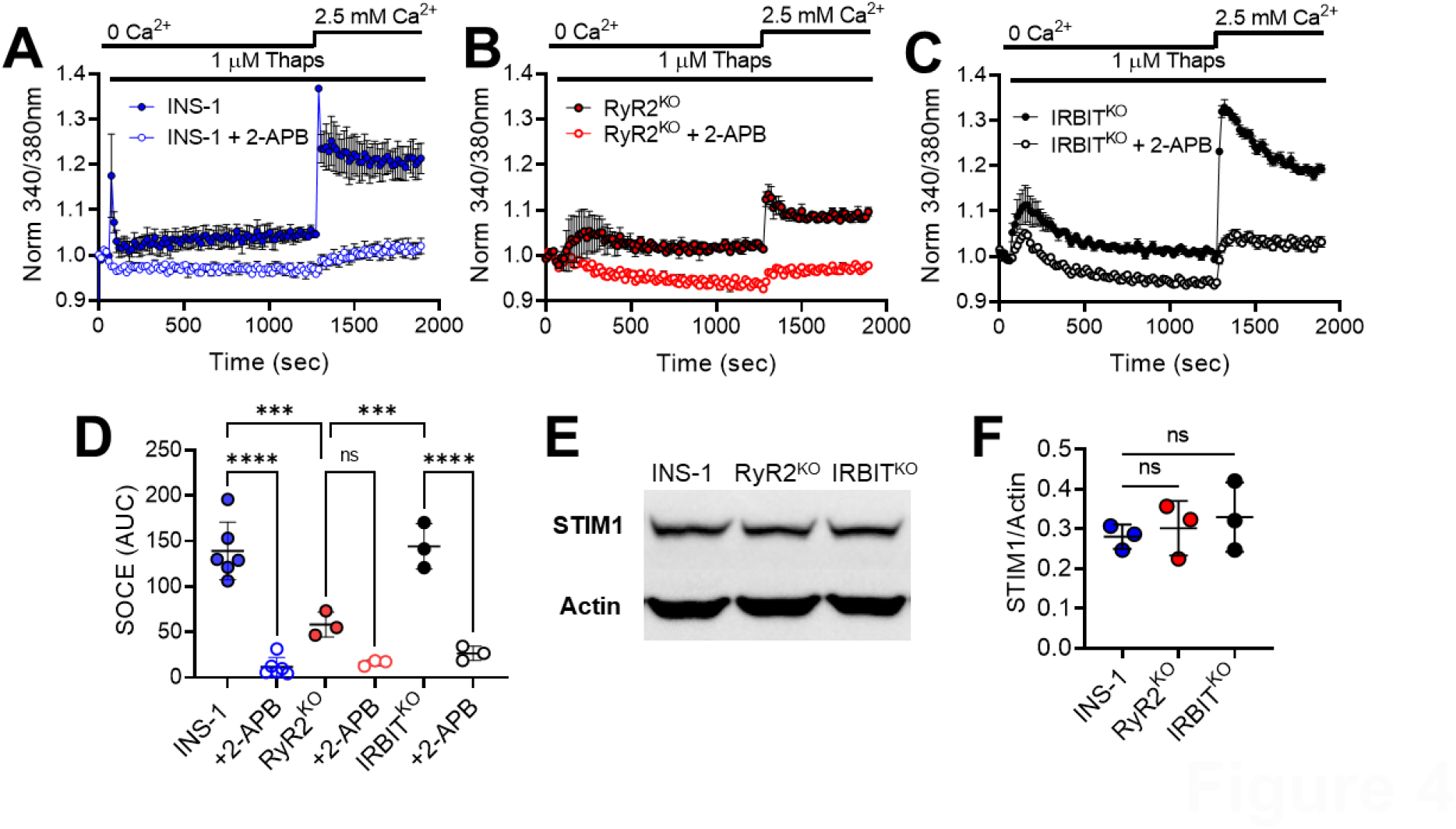
SOCE is diminished in the absence of RYR2, but not IRBIT-. Representative experiments showing activation of SOCE in control INS-1 cells **A)**, in RyR2^KO^ cells **B)**, and in IRBIT^KO^ cells **C)**. ER Ca^2+^ stores were depleted with thapsigargin in the absence of extracellular Ca^2+^, and SOCE initiated by increasing extracellular Ca^2+^ to 2.5 mM. Each point is the mean of three replicates and is shown ± SE. **D)** Quantification of SOCE (AUC). The SOCE Ca^2+^ integral in significantly reduced in RyR2^KO^ cells compared to either control INS-1 cells or IRBIT^KO^ cells (***, *P* < 0.001). 2-APB (100 µM) significantly reduced SOCE in control INS-1 cells and IRBIT^KO^ cells, but not in RyR2^KO^ cells (****, *P* < 0.0001). Two-way ANOVA with Tukey’s multiple comparisons test. INS-1 cell: n = 6 separate experiments; INS-1 cells + 2-APB: n = 6 separate experiments; RyR2^KO^ cells: n = 3 separate experiments; RyR2^KO^ cells + 2-APB: n = 3 separate experiments. **E)** Immunoblot for STIM1 from cell lysates prepared from control INS-1, RyR2^KO^, and IRBIT^KO^ cells. Representative of three separate experiments. **F)** Quantification of STIM1 immunoblots from INS-1, RyR2^KO^, and IRBIT^KO^ cell lysates. The intensity of STIM1 bands was normalized to that of actin in each replicate (n = 3 for all cell lines). No significant difference in the STIM1/actin ratios among the three cell lines was detected (One-way ANOVA).

### 3.4 Voltage-gated calcium channel current density is elevated in RyR2^KO^cells

PIP_2_ is known to be an essential regulator of channel function at the plasma membrane, including regulation of voltage-gated calcium channels [37, 38]. Given that RyR2^KO^ cells have reduced basal and stimulated PLC activity, and IRBIT^KO^ and RyR2^KO^ cells have increased PIP_2_ levels, we examined the Ca_v_ current density in RyR2^KO^, IRBIT^KO^, and control INS-1 cells using whole-cell voltage clamp. Current traces were recorded from all three cell lines using Ba^2+^ as the permeant ion (**Fig. 5A**). Plots of the current density (pA/pF) voltage relationship for each cell type are summarized in **Fig. 5B**. Peak current density was significantly greater for both RyR2^KO^ and IRBIT^KO^ cells compared to control INS-1 cells, and was significantly greater in RyR2^KO^ cells compared IRBIT^KO^ cells (**Fig. 5C**). This increased peak current density in RyR2^KO^ and IRBIT^KO^ cells versus control cells was observed in the presence and absence of perfusion with extracellular solution, and in a solution set with no extracellular NaCl (**Fig. S2**). Further, the increase in peak current density in RyR2^KO^ cells versus control cells was also observed when Ca^2+^, rather than Ba^2+^, was used as the permeant cation (**Fig S2**). To determine if upregulation of a particular family of Ca_v_ channels might be responsible for this upregulation current density, we examined the voltage-dependence of activation of Ca_v_ current from all three cell lines, and found no difference between either RyR2^KO^ cells or IRBIT^KO^ cells and controls cells (**Fig. 5 D**). To determine the percentage of current contributed by L-type channels (Ca_v_1.2 and Ca_v_1.3), we measured the fraction of current blocked by a maximally effective concentration (5 µM) of the L-type channel selective inhibitor, nifedipine. The fraction of current blocked by nifedipine in control INS-1 cells was ∼20%, and was not significantly different in RyR2^KO^ or IRBIT^KO^ cells (**Fig. 5E**). These results suggest a general upregulation of Ca_v_ channel activity in RyR2^KO^ and IRBIT^KO^ cells, since the fraction of current contributed by L-type channels and non-L-type channels remains unchanged.

**Figure 5.**
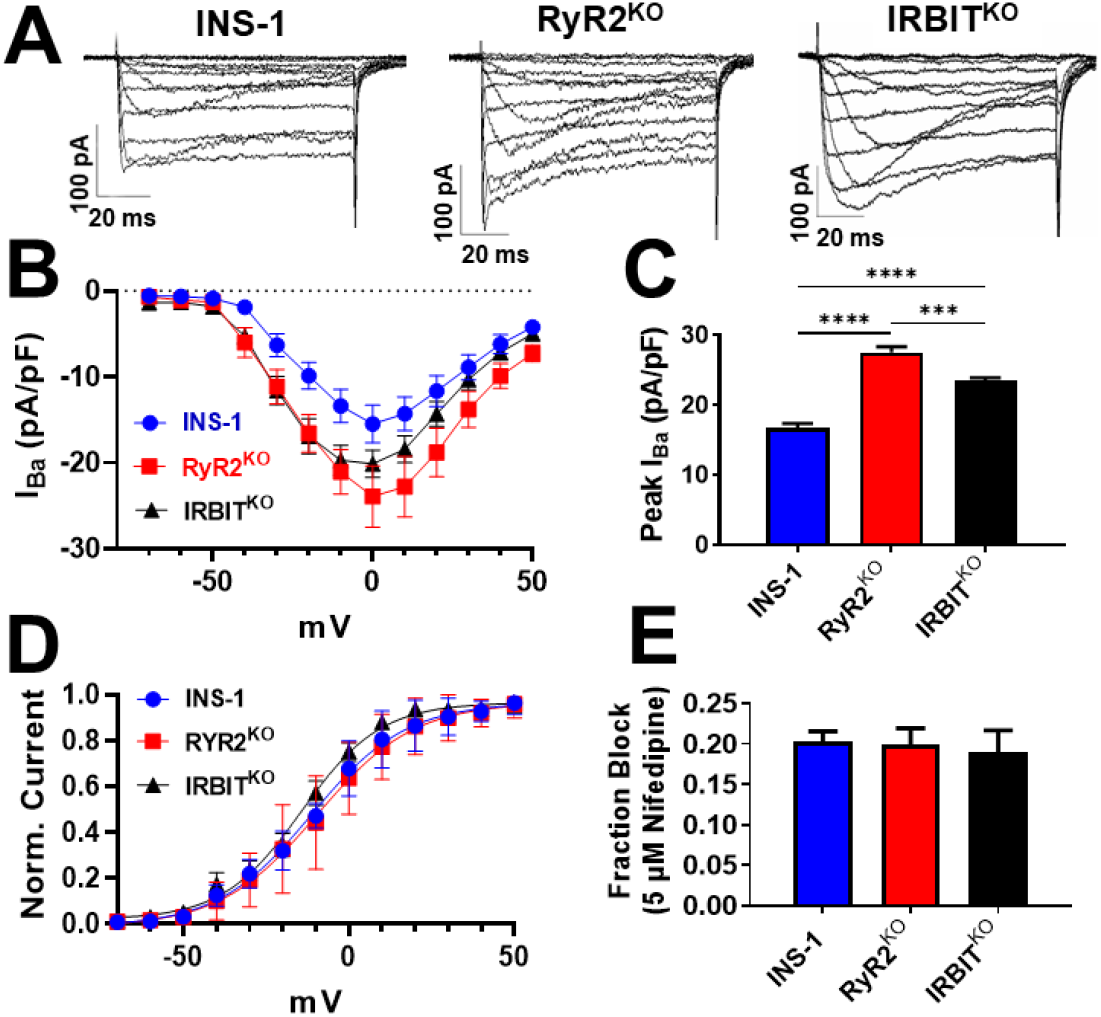
Deletion of RyR2 or IRBIT increases Ca_v_ channel activity-. **A)** Example current ensembles, recorded from the indicated cell types, elicited by stepping to voltages ranging from -70 V to +50 V for 100 s from a holding potential of -80 V. **B)** Current-voltage relationship of complied data from the indicated cell types. Data are shown as mean ± SE. **C)** Peak Barium current (I_Ba_) density of RyR2^KO^ and IRBIT^KO^ cells was greater than that of control INS-1 cells (****, *P* < 0.0001) and greater in RyR2^KO^ cells than in IRBIT^KO^ cells (***, *P* < 0.001; One-way ANOVA with Tukey’s multiple comparisons test). **D)** Voltage-dependence of activation (V_1/2_ act.) was not different between INS-1 (−9.3 ± 1.2 mV; n = 11) and RyR2^KO^ (−7.5 ± 1.0 mV; n = 24), or IRBIT^KO^ cells (−13.7 ± 1.2 mV; n = 19) (One-way ANOVA). **E)** The fraction of current blocked by 5 µM nifedipine wasn’t different between INS-1, RyR2^KO^, and IRBIT^KO^ cells (One-way ANOVA). INS-1 cells: n = 11; RyR2^KO^ cells: n = 24; IRBIT^KO^ cells: n = 19.

### 3.5 Hydrolysis of PIP_2_ preferentially reduced current in RyR2^KO^ cells

Given the increased Ca_v_ current density measured in both RyR2^KO^ and IRBIT^KO^ cells, we sought to identify a likely mechanism to explain it. We concentrated on the RyR2^KO^ cells line in these studies, since it had the greatest increase in Ca_v_ current density compared to control cells. PIP_2_ is a key regulator of actin polymerization [39], and cortical actin filaments (f-actin) are known to positively regulate voltage-gated calcium channel activity [40]. Therefore, control INS-1 and RyR2^KO^ cells were fixed, stained with phalloidin-CF405, and imaged with confocal microscopy to measure relative amounts of cortical f-actin (**Fig. 6A**). Surprisingly, RyR2^KO^ cells had ∼80% lower intensity of cortical f-actin staining compared to control INS-1 cells (**Fig. 6B**). To further assess the contribution of PIP_2_ to the increase in Ca_v_ current density, RyR2^KO^ and control cells were transfected with 2 plasmids encoding a rapamycin-inducible phosphatase system (**Fig. 6C)**. In the presence of rapamycin, the lipid phosphatase pseudojanin (PJ), fused to FKBP, localizes to the plasma via a membrane-anchored co-receptor (FRB) for rapamycin, and rapidly depletes PIP_2_ [31]. Application of 1 μM rapamycin to RyR2^KO^ cells expressing these plasmids reduced current amplitude in RyR2^KO^ cells to a greater extent than in control INS-1 cells. Application of rapamycin to cells expressing pseudojanin lacking phosphatase activity (PJ-Dead, or PJ-D) failed to reduce current density in either cell line (**Fig. 5D**). Thus, deletion of RyR2 results in greatly reduced levels of cortical f-actin, but this doesn’t likely explain the increased Ca_v_ current density in RyR2^KO^ cells compared to control INS-1 cells. However, hydrolysis of PIP_2_ rapidly reduces Ca_v_ current in RyR2^KO^ preferentially over controls INS-1 cells.

**Figure 6.**
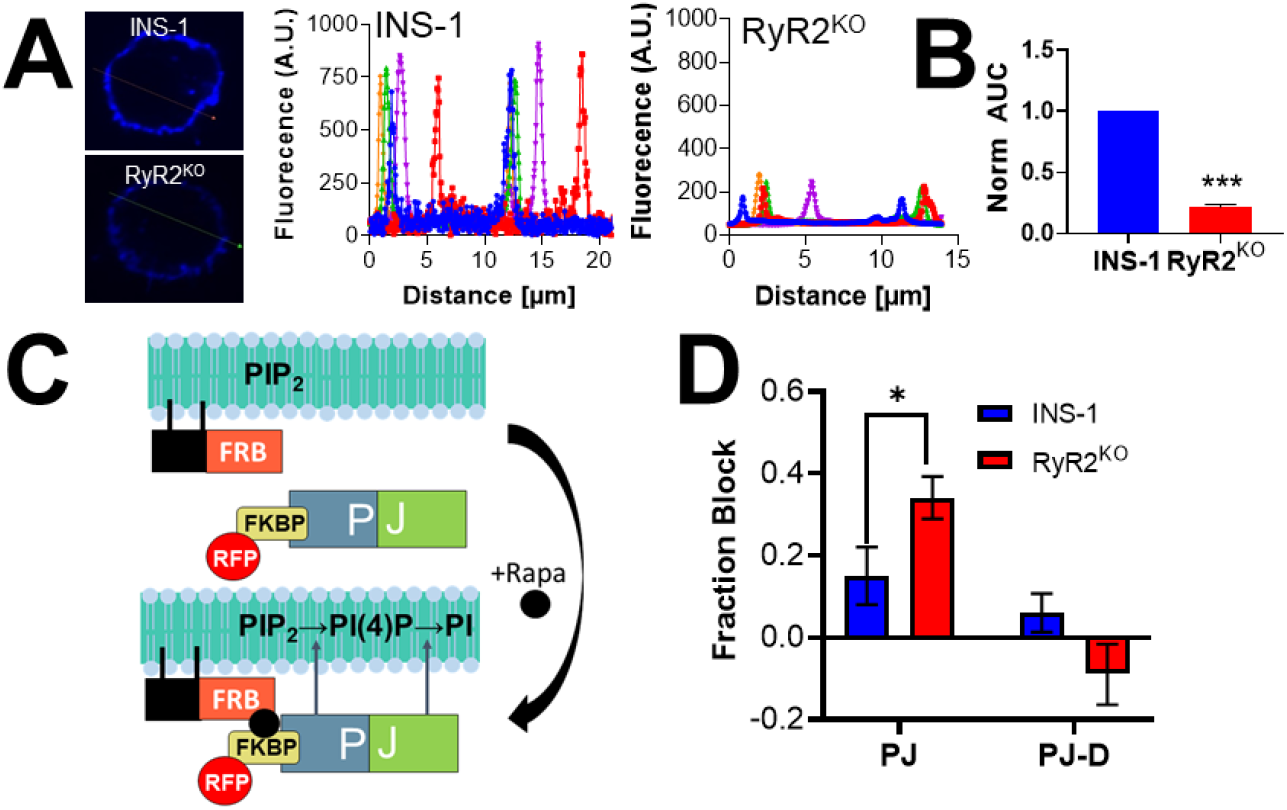
Increased plasma membrane PIP_2_ levels contributes to the increased Ca_v_ channel activity in RyR2^KO^ cells-. **A)** Cortical f-actin levels are reduced in RyR2^KO^ cells compared to controls. Fluorescence intensity of phallodin-CF405 was detected by confocal microscopy, and quantified using line scans of stained cells. **B)** Comparison of phalloidin-CF405 fluorescence intensity in RyR2^KO^ cells and INS-1 cells, normalized to INS-1 cells. Cortical CF405 fluorescence was significantly lower in RyR2^KO^ cells compared to control INS-1 cells (***, *P* < 0.001; un-paired t-test). INS-1 cells: n = 10; RyR2^KO^ cells: n = 10. **C)** Diagram of the pseudojanin (PJ) lipid phosphatase system [31]. Rapamycin induces dimerization of FK506 binding protein 12 (FKBP), and fragment of mTOR that binds rapamycin (FRB). FRB is fused to the plasma membrane-localizing peptide lyn_11_, and dimerization drives FKBP-PJ fusion to the plasma membrane. PJ thus localized, rapidly degrades PIP_2_ in the plasma membranes. Cells expressing these constructs can be identified by detection of red fluorescent protein (RFP) emission. **D)** Rapamycin perfusion (1 µM) reduces current to a greater extent in RyR2^KO^ cells expressing PJ than in INS-1 cells expressing PJ (*, *P* < 0.05; unpaired t-test). Rapamycin perfusion has no effect on current in cells expressing the phosphatase-dead PJ construct PJ-D. INS-1 cells + PJ: n = 13; INS-1 cells +PJ-D: n = 8; RyR2^KO^ cells + PJ: n = 14; RyR2^KO^ cells + PJ-D: n = 4.

### 3.6 Deletion of RYR2 increases action potential frequency

Given the changes in Ca_v_ channel current density observed in RyR2^KO^ cells, we examined the excitability of both control INS-1 cells and RyR2^KO^ cells by measuring changes in membrane potential in response to elevated glucose concentrations using current clamp in the zero-current injection (I = 0) mode. To elicit depolarization and firing of action potentials (AP), cells under current clamp were perfused with a solution containing 2.5 mM glucose to establish a baseline membrane potential, then switch to a solution containing 18 mM glucose (**Fig. 7A**). Average action potential frequencies were determined using uniform bursts of action potentials and excluding gaps between bursts. We found that the glucose-stimulated action potential frequency is doubled by RyR2 deletion. Control INS-1 cells displayed a mean action potential frequency of 0.94 Hz while that in RyR2^KO^ cell was 2.16 Hz (**Fig 7B**). Moreover, when 1 µM apamin, a blocker of the SK (KCa2) channel) [41]., was applied to control INS-1 cells during a train of action potentials, the firing rate increased to 2.18 Hz (**Fig. 7A and B**). In contrast, application of 1 µM apamin to RyR2^KO^ cells during a train of action potentials had no effect on firing frequency (**Fig 7A and B**). Then main mechanism whereby apamin enhances firing frequency in β-cells is by reducing the SK channel-mediated afterhyperpolarization (AHP) at the end of each action potential [5]. Therefore, we measured the AHP amplitudes in both control INS-1 and RyR2^KO^ cells by measuring the difference between the lowest potential and the previous plateau potential. The AHP amplitudes measured for INS-1 and RyR2^KO^ cells were: -11.52 mV and -5.98 mV, respectively in the absence of 1μM apamin, -5.38 mV and -6.96 mV, respectively, in the presence of 1μM apamin (**Fig. 7C**). Thus, the increased glucose-stimulated action potential frequency and reduced AHP amplitude in RyR2^KO^ cells, along with their unresponsiveness to apamin, argue that SK channels aren’t activated during glucose stimulation in RyR2^KO^ cells (**Fig. 7D**).

**Figure 7.**
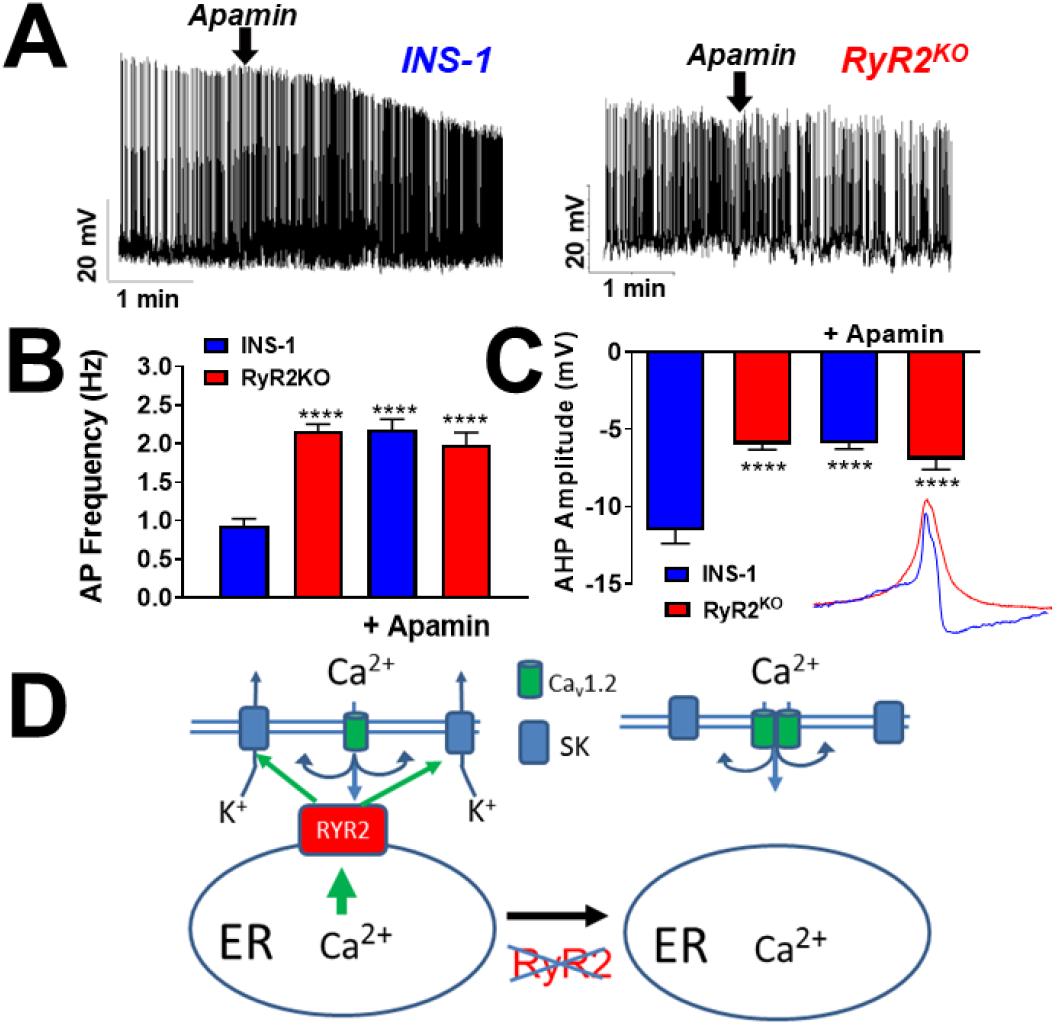
Deletion of RyR2 inhibits activation of SK channels during glucose stimulated electrical activity-. **A)** Example trains of glucose-stimulated (18 mM) action potentials in INS-1 and RyR2^KO^ cells. The SK channels blocker apamin (1 µM) was added at the time indicated by the arrows. **B)** Glucose-stimulated action potentials were more frequent in RyR2^KO^ cells than in control INS-1 cells. Addition of apamin increased action potential frequency 2-fold in control INS-1 cells, but had no effect on action potential frequency in RyR2^KO^ cells. (****, P < 0.0001; One-way ANOVA with Tukey’s multiple comparisons test) INS-1 control: n = 9 cells; RyR2^KO^ control: n = 15 cells; INS-1 + apamin: n = 10 cells; RyR2^KO^ + apamin: n = 7 cells **C)** Afterhyperpolarization (AHP) amplitude of glucose-stimulated action potentials is reduced in RyR2^KO^ cells compared to control INS-1 cells. Apamin reduces AHP amplitude in control INS-1 cells, but not in RyR2^KO^ cells. (****, *P* < 0.0001; One-way ANOVA with Tukey’s multiple comparisons test). Inset-overlay of action potentials from a control INS-1 cells (blue) and an RyR2^KO^ cells (red) illustrates the difference AHP amplitude. INS-1 control: 8,698 action potentials from 8 cells; RyR2^KO^ control: 21,241 action potentials from 8 cells; INS-1 + apamin: 5371 action potentials from 6 cells; RyR2^KO^ + apamin: 7018 action potentials from 6 cells. **D)** Model for RyR2 control of glucose-stimulated action potential frequency. In control cells, Ca^2+^ release from RyR2 induced by Ca^2+^ influx from voltage-gated Ca^2+^ channels actives SK channels during glucose stimulation. In the absence of RyR2, the enhanced Ca^2+^ influx (i.e. increased Ca_v_ channel current density and increased action potential frequency) isn’t able to activate SK channels. However, the reduced SOCE observed in RyR2^KO^ cell could also account for the deficit in SK channel activation in these cells.

## 4. Discussion

### 4.1 Role of RyR2 in Pancreatic β-cells

The role of RyRs in pancreatic β-cells has been the subject of much debate. Evidence for CICR stimulated by Ca^2+^ influx via Ca_v_ channels was reported over twenty years ago [3], and various roles for RyRs in β-cells have been reported [9, 42, 43]. Genetic studies in humans revealed that patients with mutations in RyR2 associated with severe arrhythmias were also prone to glucose intolerance [44]. In addition, knock-in of these gain of function mutations in mice also resulted in glucose intolerance [44]. Recently, RyR2 protein was detected in both INS-1 cells and murine β-cells using the highly sensitive and specific targeted mass spectrometry technique [10], and RyR2 was determined to be the major RyR transcript present in highly purified human β-cells [8]. Peptides derived from the intracellular II-III loop of Ca_v_1.2 modulate RyR2 activity [45], and expression of a peptide corresponding to the entire II-III loop of Ca_v_1.2 disrupts CICR in INS-1 cells [5]. Previously, we found that deletion of RyR2 from INS-1 cells abolishes the Ca^2+^ transient stimulated by caffeine, suggesting that RyR2 is the major functional RyR in these cells [11]. RyR2 deletion from INS-1 cells also strongly reduced insulin content, transcript, and basal and glucose stimulated secretion, and markedly reduced levels of the protein IRBIT [11]. In this study, we examined the role of RyR2 and IRBIT in regulation of key Ca^2+^ signaling processes in INS-1 cells.

### 4.2 RyR2 regulates Tolbutamide-stimulated Ca^2+^ transients and insulin secretion

Depolarization-dependent Ca^2+^ influx is a key driver of insulin secretion [46]. Tolbutamide stimulates a biphasic Ca^2+^ response, with a rapid peak, followed by sustained plateau of elevated [Ca^2+^]_in_ (**Fig 1A**). This rapid peak was inhibited by ryanodine (**Fig. 1B**) [47] and by a peptide derived from an intracellular loop of Ca_v_1.2 [5]. RyR2 deletion similarly abolished the rapid peak in [Ca^2+^]_in_ in response to stimulation by tolbutamide as assessed by a plate-based assay with a short time window (2 min) (**Fig. 1B**). However, in single-cell experiments with a longer time frame (5 min), tolbutamide stimulated marked peaks in [Ca^2+^]_in_ in both RyR2^KO^ and IRBIT^KO^ cells that were delayed (by 110 sec and 53 sec, respectively) compared to the rapid peak observed in control INS-1 cells. This delay corresponds with the contribution of IP_3_R to the Ca^2+^ transient in RyR2^KO^ and IRBIT^KO^ cells but not in control INS-1 cells (**Fig. 1E**). Even though the Ca^2+^ integral in response to tolbutamide wasn’t reduced by RyR2 deletion, basal and tolbutamide-stimulated insulin secretion were reduced 53 and 64%, respectively, compared to control INS-1 cells **(Fig. 1F**). This deficit in secretion observed in RyR2^KO^ cells persisted upon co-stimulation with an EPAC-selective cAMP analog and tolbutamide (42% reduction compared to control INS-1 cells). These deficits in tolbutamide-stimulated insulin secretion mirror those reported for glucose-stimulated insulin secretion in RyR2^KO^ cells [11]. Since tolbutamide directly inhibits the K_ATP_ channel [48], it’s unlikely that deficits in glucose-stimulated insulin secretion in RyR2^KO^ cells reflect a deficit in glucose metabolism, but result from the marked reduction in insulin transcript and content in these cells [11]. This conclusion is also supported by the observation that stimulation of electrical activity by glucose in RyR2^KO^ cells is intact, with a higher firing rate than control INS-1 cells (**Fig. 7**). Similarly, the reduction in tolbutamide-stimulated insulin secretion in IRBIT^KO^ cells mirrors the deficit in glucose-stimulated insulin secretion reported for these cells [11]. Given that the Ca^2+^ integral upon tolbutamide stimulation in IRBIT^KO^ cells isn’t different (**Fig. 1E**), and the Ca_v_ channel current density was increased compared to control INS-cells (**Fig. 6**), it’s also likely that the deficits in insulin secretion stimulated by tolbutamide (**Fig. 1F**) or glucose in IRBIT^KO^ cells results from decreased insulin transcript and content [11]. However, even though an EPAC-selective cAMP analog significantly potentiated tolbutamide-stimulated insulin secretion in all three cells lines (**Fig. 1F**), and activation of EPAC2 strongly potentiates stimulated insulin granule transport to the plasma membrane and fusion [49], a deficit in granule trafficking upon RyR2 or IRBIT deletion can’t be ruled out.

### 4.3 RYR2 regulates SOCE and PLC activity

The deletion of RyR2 and IRBIT both permitted activation of IP_3_R receptors during tolbutamide stimulation (**Fig. 1E**), and increased the effectiveness of 50 µM bethanechol in stimulating increases in [Ca^2+^]_in_ compared to control INS-1 cells (**Fig. 2B**). Given this, we examined the activation of PLC by the muscarinic receptor agonist carbachol. A positive feedback mechanism whereby Ca^2+^ release via IP_3_R directly, or indirectly via depletion of ER Ca^2+^ and activation of SOCE, is proposed to support sustained PLC activity [26]. Our finding that PLC activation in response to glucose is reduced in both RyR2^KO^ and IRBIT^KO^ cells, with a significantly greater deficit in RyR2^KO^ cells **(Fig. 2C**), suggests that under these conditions, the reduced PLC activation mirrors deficits in secretion. It’s possible insulin [50] or ATP [51] secretion by β-cells has an autocrine effect via activation of PLC. It will be of interest to determine if either of these mechanisms can account for this observation. In contrast, the stimulation of PLC by carbachol was sharply reduced in RyR2^KO^ cells, but not different from controls in IRBIT^KO^ cells as assessed with the IP_1_ accumulation assay (**Fig. 2C**). The marked effect of RyR2 deletion on carbachol-stimulated PLC activity was further corroborated using TIRFm imaging of GFP-C1-PLCdelta-PH at the plasma membrane (**Fig. 3C-E**). In both assays, the SOCE inhibitor 2-APB strongly inhibited PLC activation in control INS-1 cells, so we examined SOCE stimulated by the sarco/endoplasmic reticulum Ca^2+^-ATPase inhibitor thapsigargin. The marked inhibition of SOCE specifically in RyR2^KO^ cells suggests that SOCE plays a role in maintaining PLC activity in response to muscarinic receptor activation, in agreement with Thore et al. [26]. Moreover, these results suggest that RyR2 plays a key role in maintaining SOCE, as proposed by Lin et al. [24], via a mechanism that doesn’t involve regulation of STIM1 protein levels (**Fig. 4 E, F**). However, it is not clear from our studies how or if RyR2 deletion interferes with functional coupling of STIM1 to SOCE. The decrease in SOCE activity could potentially account for the sharply reduced basal insulin secretion in RyR2^KO^ cells, since SOCE is increased during ER stress leading to enhanced basal insulin secretion [52].

### 4.4 RyR2 regulates PIP_2_ levels and Ca_v_ channel current

Influx of Ca^2+^ via voltage-gated Ca^2+^ channels is a key regulator of insulin secretion since inhibitors of both L-type [53] [4] and non-L-type [4] channels are able to substantially inhibit secretion. Our finding that Ca_v_ channel current density is upregulated in RyR2^KO^ and IRBIT^KO^ cells (**Fig. 5A-C**) contrasts with our finding that insulin secretion in these cell lines is reduced (**Fig 1F)**. The lack of any significant change in activation potential (**Fig. 5D**), or in the fraction of current blocked by the L-type channel inhibitor nifedipine (**Fig. 5E**), suggest a general increase in Ca_v_ channel activity. However, it’s possible that the complement of L-type (i.e. Ca_v_1.2, Ca_v_1.3) or non-L-type channel subtypes (Ca_v_2.2, Ca_v_2.3) has changed without changing the ratio of nifedipine-sensitive to nifedipine-insensitive current. A prominent role of plasma membrane PIP_2_ levels in this upregulation of Ca_v_ current is supported by the experiments with the rapamycin-activated phospholipid phosphatase pseudojanin [31], showing a greater decrease in current upon depletion of PIP_2_ in RyR2^KO^ cells than in control INS-1 cells. We propose that the increase in Ca_v_ channel current density is a result of increased PIP_2_ levels at the plasma membrane (**Fig. 3A, B**). PIP_2_ is potentiator of Ca_v_ channels activity in general [54], and specifically in pancreatic β-cells [28]. In the case of RyR2^KO^ cells, the increased PIP_2_ levels could follow from sharply reduced PLC activity (**Fig. 2B and 3E**). Deletion of IRBIT leads to an increase in PIP_2_ levels as well, and this could be ascribed to the decreased basal and glucose-stimulated PLC activity observed in this cell line (**Fig 2B**). However, IRBIT also binds to the catalytic core of phosphatidylinositol phosphate kinases (PIPK) type Iα and type IIα [55], key enzymes in the production of PIP_2_. This binding is competitive with Mg^2+^, ATP, and phosphatidylinositol-4-phosphate, but had no discernable effect on kinase activity *in vitro*. We speculate that, in cells, IRBIT binding may negatively regulate PIPK such that deletion or reduction of IRBIT enhances plasma membrane PIP_2_ levels, and contributes to the observed increase in Ca_v_ channel current density.

### 4.5 RyR2 regulates glucose-stimulated electrical activity

Glucose-stimulated action potentials in pancreatic β-cells are mediated by the membrane depolarizing effect of K_ATP_ channel closure upon ATP binding. As the membrane potential is depolarized, Ca_v_ channels are activated, leading to the upstroke of the action potential, followed by activation of K_v_ channels which repolarize the membrane potential [56]. This process is fine-tuned by small conductance, Ca^2+^-activated K^+^ (SK) channels (K_Ca_2.1-3) [41]. Activation of SK channels regulates action potential frequency by conducting a Ca^2+^-dependent activation of K^+^ efflux late in the repolarization phase, referred to as the afterhyperpolarization (AHP) [57]. The magnitude of the AHP controls AP frequency because it largely dictates the time required for spontaneous depolarization to reach the threshold for firing. We previously demonstrated that overexpression of the Cav1.2 II-III intracellular loop in INS-1 cells increases action potential frequency through disruption of CICR and SK channel activity [5]. In this study, we found that deletion of RyR2 from INS-1 cells has the same effect on glucose-stimulated action potential frequency. In RyR2^KO^ cells, glucose stimulated action potentials with a mean frequency of 2.2 Hz, and these action potentials were in sensitive to the SK channel blocker apamin (**Fig. 7B**). In contrast, the mean frequency in control INS-1 cells was 0.94 Hz, which was increased to 2.3 Hz in the presence of apamin. Similarly, the AHP amplitude in RyR2^KO^ cells was -6.0 mV in the absence of apamin, and -7.0 mV in the presence of apamin. In contrast, the AHP amplitude in the absence of apamin in control INS-1 cells was -11.5 mV, and -5.9 mV in the presence of apamin (**Fig. 7C**). Thus, both the frequency of action potentials and the amplitude of the AHP in RyR2^KO^ cells in the absence of apamin were indistinguishable from those in the presence of apamin in both RyR2^KO^ and control cells, indicating a failure in SK channel activation in RyR2^KO^ cells. As expected, apamin significantly increased action potential frequency and decreased AHP amplitude in control INS-1 cells. These results, along with the observation that RyR2^KO^ cells have greater Ca_v_ channel current density and release of Ca^2+^ from IP_3_ receptors than controls, suggest that Ca^2+^ release from RyR2 plays a unique role in activating SK channels in INS-1 cells (**Fig. 7D**). However, the reduced SOCE observed in RyR2^KO^ cells could also account for the deficit in SK channel activation in these cells.

## 5 Conclusions

This study examined the role of RyR2 in several aspects of Ca^2+^ signaling in the pancreatic β-cell line INS-1. The major findings (Summarized in **Fig. 8**) are that 1: Deletion of RyR2 delays the major peak in [Ca^2+^]_in_ in response to tolbutamide stimulation, and enhances release of Ca^2+^ IP_3_ receptors; 2: Deletion of RyR2 markedly reduces basal insulin secretion and insulin secretion in response to tolbutamide stimulation; 3: Deletion of RyR2 markedly reduces basal and stimulated PLC activity; 4: Deletion of RyR2 reduces SOCE in response to emptying of ER Ca^2+^ stores with thapsigargin; 5: Deletion of RyR2 increases Ca_v_ channel current density via an increase in plasma membrane PIP2 levels; 6: Deletion of RyR2 inhibits activation of SK channels and increases glucose-stimulated action potential frequency. Thus, via regulation of SOCE activity, RyR2 controls PLC activity and likely PIP_2_ levels. It will be of interest to determine the molecular mechanism that links RyR2 to SOCE.

**Figure 8.**
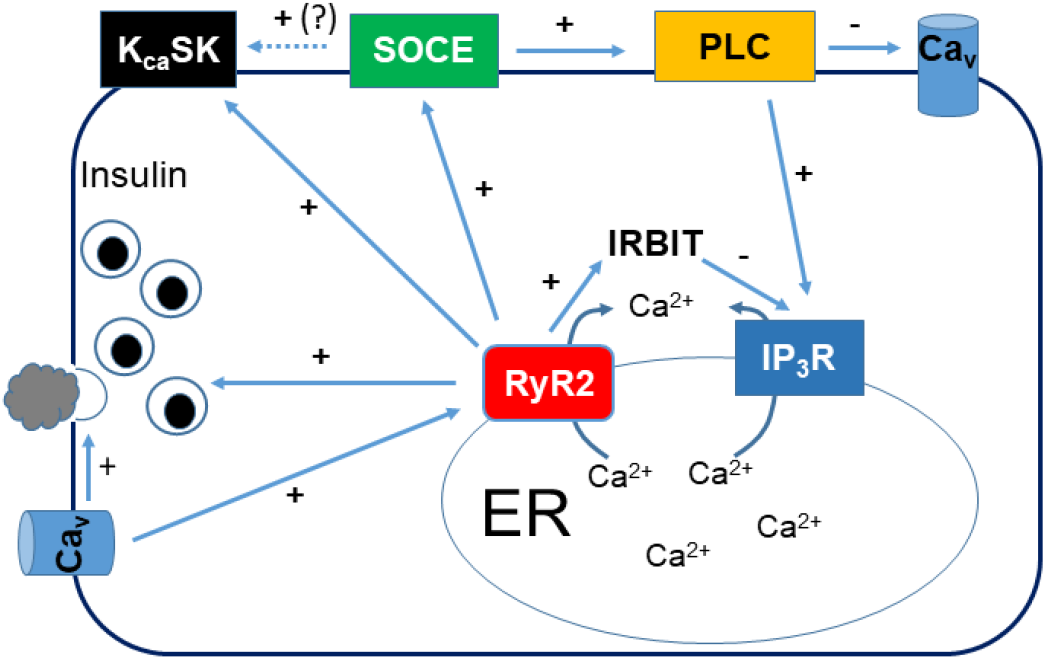
Model for regulation of Ca^2+^ dynamics in pancreatic β-cells-. RyR2 is regulated by influx of Ca^2+^ via L-type Ca_v_ channels [3, 5], which can also directly stimulate rapid insulin release [58]. RyR2 regulates insulin secretion either directly via Ca^2+^ release from the ER or indirectly via regulation of insulin content and transcript level [11]. RyR2 plays a key role in regulating SOCE via a yet unknown mechanism. Maintenance of SOCE is critical for basal and stimulated PLC activity. Reduced PLC activity in the absence of RyR2 increases plasma membrane PIP_2_ levels, and increases Ca_v_ channel activity. RyR2 also directly or indirectly (via regulation of SOCE) regulates activation of SK channels. Finally, RyR2 regulates IP_3_R-mediated Ca^2+^ release, and perhaps many other cellular processes, via regulation of IRBIT levels [11].

## Abbreviations

Ca_v_: Voltage-gated Ca^2+^
CICR: Ca^2+^-induced Ca^2+^ release
EPAC: Exchange protein directly activated by cyclic AMP
ER: Endoplasmic reticulum
IRBIT: IP_3_ receptor binding protein released with inositol 1,4,5-trisphosphate
IP_3_: Inositol trisphosphate
IP_3_R: Inositol trisphosphate receptor
PIP_2_: Phosphatidyl inositol-4,5-bisphosphate
PLC: Phospholipase C
RyR2: Ryanodine receptor 2
SOCE: Store-operated Ca^2+^ entry
STIM1: Stromal interaction molecule 1
SK: Small-conductance Ca^2+^-activated K^+^
TIRFm: Total internal reflection fluorescence microscopy

## CRediT author contribution statement

Kyle E. Harvey: Investigation, Visualization, Writing-review & editing, Formal analysis. Shiqi Tang: Investigation, Visualization, Formal analysis. Emily K. LaVigne: Investigation, Visualization, Formal analysis, Writing-review and editing. Evan P.S. Pratt: Investigation, Visualization, formal analysis. Gregory H. Hockerman: Formal analysis, Visualization, Writing original draft, Writing-Review & Editing, Supervision, Project administration, Funding Acquisition.

## Acknowledgements

The authors thank Dr. Robert Stahelin for the gift of PIP_2_ antibodies. This work was supported by a Richard and Anne Borch Award (GHH), and a Showalter Faculty Scholar Award (GHH).

## Appendix. Supplementary Materials

Figure S1

Figure S2

## Data Availability

Data will be made available on request.

## References

1. Rorsman P, Ashcroft FM. Pancreatic beta-Cell Electrical Activity and Insulin Secretion: Of Mice and Men. Physiological reviews. 2018;98(1):117–214. doi: 10.1152/physrev.00008.2017. PubMed PMID: 29212789; PubMed Central PMCID: PMCPMC5866358.

2. Sabatini PV, Speckmann T, Lynn FC. Friend and foe: beta-cell Ca(2+) signaling and the development of diabetes. Molecular metabolism. 2019;21:1–12. Epub 2019/01/12. doi: 10.1016/j.molmet.2018.12.007. PubMed PMID: 30630689; PubMed Central PMCID: PMCPMC6407368.

3. Lemmens R, Larsson O, Berggren PO, Islam MS. Ca2+-induced Ca2+ release from the endoplasmic reticulum amplifies the Ca2+ signal mediated by activation of voltage-gated L-type Ca2+ channels in pancreatic beta-cells. The Journal of biological chemistry. 2001;276(13):9971–7. doi: 10.1074/jbc.M009463200. PubMed PMID: 11139580.

4. Braun M, Ramracheya R, Bengtsson M, Zhang Q, Karanauskaite J, Partridge C, et al. Voltage-gated ion channels in human pancreatic beta-cells: electrophysiological characterization and role in insulin secretion. Diabetes. 2008;57(6):1618–28. doi: 10.2337/db07-0991. PubMed PMID: 18390794.

5. Wang Y, Jarrard RE, Pratt EP, Guerra ML, Salyer AE, Lange AM, et al. Uncoupling of Cav1.2 from Ca(2+)-induced Ca(2+) release and SK channel regulation in pancreatic beta-cells. Molecular endocrinology. 2014;28(4):458–76. doi: 10.1210/me.2013-1094. PubMed PMID: 24506535; PubMed Central PMCID: PMCPMC3968403.

6. Arnette D, Gibson TB, Lawrence MC, January B, Khoo S, McGlynn K, et al. Regulation of ERK1 and ERK2 by glucose and peptide hormones in pancreatic beta cells. The Journal of biological chemistry. 2003;278(35):32517–25. doi: 10.1074/jbc.M301174200. PubMed PMID: 12783880.

7. Bruton JD, Lemmens R, Shi CL, Persson-Sjogren S, Westerblad H, Ahmed M, et al. Ryanodine receptors of pancreatic beta-cells mediate a distinct context-dependent signal for insulin secretion. FASEB journal: official publication of the Federation of American Societies for Experimental Biology. 2003;17(2):301–3. doi: 10.1096/fj.02-0481fje. PubMed PMID: 12475892.

8. Nordenskjold F, Andersson B, Islam MS. Expression of the Inositol 1,4,5-Trisphosphate Receptor and the Ryanodine Receptor Ca(2+)-Release Channels in the Beta-Cells and Alpha-Cells of the Human Islets of Langerhans. Advances in experimental medicine and biology. 2020;1131:271–9. doi: 10.1007/978-3-030-12457-1_11. PubMed PMID: 31646514.

9. Johnson JD, Kuang S, Misler S, Polonsky KS. Ryanodine receptors in human pancreatic beta cells: localization and effects on insulin secretion. FASEB journal: official publication of the Federation of American Societies for Experimental Biology. 2004;18(7):878–80. doi: 10.1096/fj.03-1280fje. PubMed PMID: 15033925.

10. Yamamoto WR, Bone RN, Sohn P, Syed F, Reissaus CA, Mosley AL, et al. Endoplasmic reticulum stress alters ryanodine receptor function in the murine pancreatic beta cell. The Journal of biological chemistry. 2019;294(1):168–81. doi: 10.1074/jbc.RA118.005683. PubMed PMID: 30420428; PubMed Central PMCID: PMCPMC6322901.

11. Harvey KE, LaVigne EK, Dar MS, Salyer AE, Pratt EPS, Sample PA, et al. RyR2/IRBIT regulates insulin gene transcript, insulin content, and secretion in the insulinoma cell line INS-1. Scientific reports. 2022;12(1):7713. doi: 10.1038/s41598-022-11276-8.

12. Lee B, Bradford PG, Laychock SG. Characterization of inositol 1,4,5-trisphosphate receptor isoform mRNA expression and regulation in rat pancreatic islets, RINm5F cells and betaHC9 cells. Journal of molecular endocrinology. 1998;21(1):31–9. Epub 1998/09/02. doi: 10.1677/jme.0.0210031. PubMed PMID: 9723861.

13. Lee B, Laychock SG. Inositol 1,4,5-trisphosphate receptor isoform expression in mouse pancreatic islets: effects of carbachol. Biochemical pharmacology. 2001;61(3):327–36. Epub 2001/02/15. doi: 10.1016/s0006-2952(00)00559-1. PubMed PMID: 11172737.

14. Luciani DS, Gwiazda KS, Yang TL, Kalynyak TB, Bychkivska Y, Frey MH, et al. Roles of IP3R and RyR Ca2+ channels in endoplasmic reticulum stress and beta-cell death. Diabetes. 2009;58(2):422–32. doi: 10.2337/db07-1762. PubMed PMID: 19033399; PubMed Central PMCID: PMC2628616.

15. Laybutt DR, Preston AM, Akerfeldt MC, Kench JG, Busch AK, Biankin AV, et al. Endoplasmic reticulum stress contributes to beta cell apoptosis in type 2 diabetes. Diabetologia. 2007;50(4):752–63. Epub 2007/02/03. doi: 10.1007/s00125-006-0590-z. PubMed PMID: 17268797.

16. Hara T, Mahadevan J, Kanekura K, Hara M, Lu S, Urano F. Calcium efflux from the endoplasmic reticulum leads to beta-cell death. Endocrinology. 2014;155(3):758–68. doi: 10.1210/en.2013-1519. PubMed PMID: 24424032; PubMed Central PMCID: PMC3929724.

17. Putney JW, Jr., Broad LM, Braun FJ, Lievremont JP, Bird GS. Mechanisms of capacitative calcium entry. Journal of cell science. 2001;114(Pt 12):2223–9. Epub 2001/08/09. doi: 10.1242/jcs.114.12.2223. PubMed PMID: 11493662.

18. Prakriya M, Lewis RS. Store-Operated Calcium Channels. Physiological reviews. 2015;95(4):1383–436. Epub 2015/09/25. doi: 10.1152/physrev.00020.2014. PubMed PMID: 26400989; PubMed Central PMCID: PMCPMC4600950.

19. Liao Y, Plummer NW, George MD, Abramowitz J, Zhu MX, Birnbaumer L. A role for Orai in TRPC-mediated Ca2+ entry suggests that a TRPC:Orai complex may mediate store and receptor operated Ca2+ entry. Proceedings of the National Academy of Sciences of the United States of America. 2009;106(9):3202–6. doi: 10.1073/pnas.0813346106. PubMed PMID: 19221033; PubMed Central PMCID: PMC2651283.

20. Venkatachalam K, van Rossum DB, Patterson RL, Ma HT, Gill DL. The cellular and molecular basis of store-operated calcium entry. Nature cell biology. 2002;4(11):E263–72. Epub 2002/11/05. doi: 10.1038/ncb1102-e263. PubMed PMID: 12415286.

21. Kono T, Tong X, Taleb S, Bone RN, Iida H, Lee CC, et al. Impaired Store-Operated Calcium Entry and STIM1 Loss Lead to Reduced Insulin Secretion and Increased Endoplasmic Reticulum Stress in the Diabetic beta-Cell. Diabetes. 2018;67(11):2293–304. doi: 10.2337/db17-1351. PubMed PMID: 30131394; PubMed Central PMCID: PMCPMC6198337.

22. Sabourin J, Le Gal L, Saurwein L, Haefliger JA, Raddatz E, Allagnat F. Store-operated Ca2+ Entry Mediated by Orai1 and TRPC1 Participates to Insulin Secretion in Rat beta-Cells. The Journal of biological chemistry. 2015;290(51):30530–9. Epub 2015/10/24. doi: 10.1074/jbc.M115.682583. PubMed PMID: 26494622; PubMed Central PMCID: PMCPMC4683273.

23. Bird GS, Putney JW, Jr. Pharmacology of Store-Operated Calcium Entry Channels. In: Kozak JA, Putney JW, Jr., editors. Calcium Entry Channels in Non-Excitable Cells. Boca Raton (FL) 2018. p. 311–24.

24. Lin AH, Sun H, Paudel O, Lin MJ, Sham JS. Conformation of ryanodine receptor-2 gates store-operated calcium entry in rat pulmonary arterial myocytes. Cardiovasc Res. 2016;111(1):94–104. Epub 2016/03/26. doi: 10.1093/cvr/cvw067. PubMed PMID: 27013634; PubMed Central PMCID: PMCPMC4909159.

25. Kadamur G, Ross EM. Mammalian phospholipase C. Annual review of physiology. 2013;75:127–54. doi: 10.1146/annurev-physiol-030212-183750. PubMed PMID: 23140367.

26. Thore S, Dyachok O, Gylfe E, Tengholm A. Feedback activation of phospholipase C via intracellular mobilization and store-operated influx of Ca2+ in insulin-secreting beta-cells. Journal of cell science. 2005;118(Pt 19):4463–71. doi: 10.1242/jcs.02577. PubMed PMID: 16159958.

27. Logothetis DE, Petrou VI, Adney SK, Mahajan R. Channelopathies linked to plasma membrane phosphoinositides. Pflugers Archiv: European journal of physiology. 2010;460(2):321–41. Epub 2010/04/17. doi: 10.1007/s00424-010-0828-y. PubMed PMID: 20396900; PubMed Central PMCID: PMCPMC4040125.

28. de la Cruz L, Puente EI, Reyes-Vaca A, Arenas I, Garduno J, Bravo-Martinez J, et al. PIP2 in pancreatic beta-cells regulates voltage-gated calcium channels by a voltage-independent pathway. American journal of physiology Cell physiology. 2016;311(4):C630–C40. Epub 2016/08/05. doi: 10.1152/ajpcell.00111.2016. PubMed PMID: 27488666.

29. Asfari M, Janjic D, Meda P, Li G, Halban PA, Wollheim CB. Establishment of 2-mercaptoethanol-dependent differentiated insulin-secreting cell lines. Endocrinology. 1992;130(1):167–78. doi: 10.1210/endo.130.1.1370150. PubMed PMID: 1370150.

30. Stauffer TP, Ahn S, Meyer T. Receptor-induced transient reduction in plasma membrane PtdIns(4,5)P2 concentration monitored in living cells. Current biology: CB. 1998;8(6):343–6. Epub 1998/03/25. doi: 10.1016/s0960-9822(98)70135-6. PubMed PMID: 9512420.

31. Hammond GR, Fischer MJ, Anderson KE, Holdich J, Koteci A, Balla T, et al. PI4P and PI(4,5)P2 are essential but independent lipid determinants of membrane identity. Science. 2012;337(6095):727–30. Epub 2012/06/23. doi: 10.1126/science.1222483. PubMed PMID: 22722250; PubMed Central PMCID: PMCPMC3646512.

32. Kang G, Joseph JW, Chepurny OG, Monaco M, Wheeler MB, Bos JL, et al. Epac-selective cAMP analog 8-pCPT-2’-O-Me-cAMP as a stimulus for Ca2+-induced Ca2+ release and exocytosis in pancreatic beta-cells. The Journal of biological chemistry. 2003;278(10):8279–85. doi: 10.1074/jbc.M211682200. PubMed PMID: 12496249; PubMed Central PMCID: PMC3516291.

33. Gilon P, Henquin JC. Mechanisms and physiological significance of the cholinergic control of pancreatic beta-cell function. Endocrine reviews. 2001;22(5):565–604. Epub 2001/10/06. doi: 10.1210/edrv.22.5.0440. PubMed PMID: 11588141.

34. Molina J, Rodriguez-Diaz R, Fachado A, Jacques-Silva MC, Berggren PO, Caicedo A. Control of insulin secretion by cholinergic signaling in the human pancreatic islet. Diabetes. 2014;63(8):2714–26. doi: 10.2337/db13-1371. PubMed PMID: 24658304; PubMed Central PMCID: PMCPMC4113066.

35. Hallcher LM, Sherman WR. The effects of lithium ion and other agents on the activity of myo-inositol-1-phosphatase from bovine brain. The Journal of biological chemistry. 1980;255(22):10896–901. Epub 1980/11/25. PubMed PMID: 6253491.

36. Bootman MD, Collins TJ, Mackenzie L, Roderick HL, Berridge MJ, Peppiatt CM. 2-aminoethoxydiphenyl borate (2-APB) is a reliable blocker of store-operated Ca2+ entry but an inconsistent inhibitor of InsP3-induced Ca2+ release. FASEB journal: official publication of the Federation of American Societies for Experimental Biology. 2002;16(10):1145–50. doi: 10.1096/fj.02-0037rev. PubMed PMID: 12153982.

37. Wu L, Bauer CS, Zhen XG, Xie C, Yang J. Dual regulation of voltage-gated calcium channels by PtdIns(4,5)P2. Nature. 2002;419(6910):947–52. Epub 2002/11/01. doi: 10.1038/nature01118. PubMed PMID: 12410316.

38. Gamper N, Reznikov V, Yamada Y, Yang J, Shapiro MS. Phosphatidylinositol [correction] 4,5-bisphosphate signals underlie receptor-specific Gq/11-mediated modulation of N-type Ca2+ channels. The Journal of neuroscience: the official journal of the Society for Neuroscience. 2004;24(48):10980–92. Epub 2004/12/03. doi: 10.1523/JNEUROSCI.3869-04.2004. PubMed PMID: 15574748; PubMed Central PMCID: PMCPMC6730206.

39. Janmey PA, Bucki R, Radhakrishnan R. Regulation of actin assembly by PI(4,5)P2 and other inositol phospholipids: An update on possible mechanisms. Biochemical and biophysical research communications. 2018;506(2):307–14. Epub 2018/08/25. doi: 10.1016/j.bbrc.2018.07.155. PubMed PMID: 30139519; PubMed Central PMCID: PMCPMC6269227.

40. Stolting G, de Oliveira RC, Guzman RE, Miranda-Laferte E, Conrad R, Jordan N, et al. Direct interaction of CaVbeta with actin up-regulates L-type calcium currents in HL-1 cardiomyocytes. The Journal of biological chemistry. 2015;290(8):4561–72. doi: 10.1074/jbc.M114.573956. PubMed PMID: 25533460; PubMed Central PMCID: PMCPMC4335199.

41. Jacobson DA, Mendez F, Thompson M, Torres J, Cochet O, Philipson LH. Calcium-activated and voltage-gated potassium channels of the pancreatic islet impart distinct and complementary roles during secretagogue induced electrical responses. The Journal of physiology. 2010;588(Pt 18):3525–37. doi: 10.1113/jphysiol.2010.190207. PubMed PMID: 20643768; PubMed Central PMCID: PMC2988516.

42. Johnson JD, Han Z, Otani K, Ye H, Zhang Y, Wu H, et al. RyR2 and calpain-10 delineate a novel apoptosis pathway in pancreatic islets. The Journal of biological chemistry. 2004;279(23):24794–802. doi: 10.1074/jbc.M401216200. PubMed PMID: 15044459.

43. Dixit SS, Wang T, Manzano EJ, Yoo S, Lee J, Chiang DY, et al. Effects of CaMKII-mediated phosphorylation of ryanodine receptor type 2 on islet calcium handling, insulin secretion, and glucose tolerance. PloS one. 2013;8(3):e58655. doi: 10.1371/journal.pone.0058655. PubMed PMID: 23516528; PubMed Central PMCID: PMC3596297.

44. Santulli G, Pagano G, Sardu C, Xie W, Reiken S, D’Ascia SL, et al. Calcium release channel RyR2 regulates insulin release and glucose homeostasis. The Journal of clinical investigation. 2015;125(5):1968–78. doi: 10.1172/JCI79273. PubMed PMID: 25844899; PubMed Central PMCID: PMC4463204.

45. Dulhunty AF, Curtis SM, Cengia L, Sakowska M, Casarotto MG. Peptide fragments of the dihydropyridine receptor can modulate cardiac ryanodine receptor channel activity and sarcoplasmic reticulum Ca2+ release. The Biochemical journal. 2004;379(Pt 1):161–72. doi: 10.1042/BJ20031096. PubMed PMID: 14678014; PubMed Central PMCID: PMCPMC1224045.

46. Devis G, Somers G, Malaisse WJ. Stimulation of insulin release by calcium. Biochemical and biophysical research communications. 1975;67(2):525-9. PubMed PMID: 812497.

47. Jarrard RE, Wang Y, Salyer AE, Pratt EP, Soderling IM, Guerra ML, et al. Potentiation of sulfonylurea action by an EPAC-selective cAMP analog in INS-1 cells: comparison of tolbutamide and gliclazide and a potential role for EPAC activation of a 2-APB-sensitive Ca2+ influx. Molecular pharmacology. 2013;83(1):191–205. doi: 10.1124/mol.112.081943. PubMed PMID: 23071106; PubMed Central PMCID: PMC3533467.

48. Cook DL, Ikeuchi M. Tolbutamide as mimic of glucose on beta-cell electrical activity. ATP-sensitive K+ channels as common pathway for both stimuli. Diabetes. 1989;38(4):416–21. Epub 1989/04/01. doi: 10.2337/diab.38.4.416. PubMed PMID: 2647550.

49. Shibasaki T, Takahashi H, Miki T, Sunaga Y, Matsumura K, Yamanaka M, et al. Essential role of Epac2/Rap1 signaling in regulation of insulin granule dynamics by cAMP. Proceedings of the National Academy of Sciences of the United States of America. 2007;104(49):19333–8. doi: 10.1073/pnas.0707054104. PubMed PMID: 18040047; PubMed Central PMCID: PMC2148290.

50. Eichhorn J, Kayali AG, Austin DA, Webster NJ. Insulin activates phospholipase C-gamma1 via a PI-3 kinase dependent mechanism in 3T3-L1 adipocytes. Biochemical and biophysical research communications. 2001;282(2):615–20. Epub 2001/06/13. doi: 10.1006/bbrc.2001.4616. PubMed PMID: 11401505.

51. Khan S, Yan-Do R, Duong E, Wu X, Bautista A, Cheley S, et al. Autocrine activation of P2Y1 receptors couples Ca (2+) influx to Ca (2+) release in human pancreatic beta cells. Diabetologia. 2014;57(12):2535–45. Epub 2014/09/12. doi: 10.1007/s00125-014-3368-8. PubMed PMID: 25208758.

52. Zhang IX, Ren J, Vadrevu S, Raghavan M, Satin LS. ER stress increases store-operated Ca(2+) entry (SOCE) and augments basal insulin secretion in pancreatic beta cells. The Journal of biological chemistry. 2020;295(17):5685–700. Epub 2020/03/18. doi: 10.1074/jbc.RA120.012721. PubMed PMID: 32179650; PubMed Central PMCID: PMCPMC7186166.

53. Liu G, Dilmac N, Hilliard N, Hockerman GH. Ca v 1.3 is preferentially coupled to glucose-stimulated insulin secretion in the pancreatic beta-cell line INS-1. The Journal of pharmacology and experimental therapeutics. 2003;305(1):271–8. doi: 10.1124/jpet.102.046334. PubMed PMID: 12649379.

54. Logothetis DE, Petrou VI, Zhang M, Mahajan R, Meng XY, Adney SK, et al. Phosphoinositide control of membrane protein function: a frontier led by studies on ion channels. Annual review of physiology. 2015;77:81–104. Epub 2014/10/09. doi: 10.1146/annurev-physiol-021113-170358. PubMed PMID: 25293526; PubMed Central PMCID: PMCPMC4485992.

55. Ando H, Hirose M, Gainche L, Kawaai K, Bonneau B, Ijuin T, et al. IRBIT Interacts with the Catalytic Core of Phosphatidylinositol Phosphate Kinase Type Ialpha and IIalpha through Conserved Catalytic Aspartate Residues. PloS one. 2015;10(10):e0141569. Epub 2015/10/29. doi: 10.1371/journal.pone.0141569. PubMed PMID: 26509711; PubMed Central PMCID: PMCPMC4624786.

56. Drews G, Krippeit-Drews P, Dufer M. Electrophysiology of islet cells. Advances in experimental medicine and biology. 2010;654:115–63. Epub 2010/03/11. doi: 10.1007/978-90-481-3271-3_7. PubMed PMID: 20217497.

57. Dwivedi D, Bhalla US. Physiology and Therapeutic Potential of SK, H, and M Medium AfterHyperPolarization Ion Channels. Front Mol Neurosci. 2021;14:658435. Epub 2021/06/22. doi: 10.3389/fnmol.2021.658435. PubMed PMID: 34149352; PubMed Central PMCID: PMCPMC8209339.

58. Barg S, Ma X, Eliasson L, Galvanovskis J, Gopel SO, Obermuller S, et al. Fast exocytosis with few Ca(2+) channels in insulin-secreting mouse pancreatic B cells. Biophysical journal. 2001;81(6):3308–23. doi: 10.1016/S0006-3495(01)75964-4. PubMed PMID: 11720994; PubMed Central PMCID: PMC1301788.

