## Supplemental Figures for "RyR2 regulates store-operated Ca^2+^ entry, phospholipase C activity, and electrical excitability in the insulinoma cell line INS-1"

**Harvey et al., Supplementary Data**
**Figure S1. Deletion of RyR2 reduces SOCE: Reduced time of exposure to 2-APB -**

Representative experiments showing activation of SOCE in control INS-1 cells **A)**, in $RyR2^{KO}$ cells **B)**, and in $IRBIT^{KO}$ cells **C)** with reduced time of exposure to 2-APB prior to re-introduction of $Ca^{2+}$**.** ER $Ca^{2+}$ stores were depleted with injection thapsigargin in the absence of extracellular $Ca^{2+}$, and SOCE initiated by increasing extracellular $Ca^{2+}$ to 2.5 mM. 100 µM 2-APB was co-injected with thapsigargin for some experiments prior to re-addition of $Ca^{2+}$ to minimize off-target effects. Each point is the mean of three replicates and is shown ± SE. **D)** Quantification of SOCE (AUC). The SOCE $Ca^{2+}$ integral in significantly reduced in $RyR2^{KO}$ cells compared to either control INS-1 cells (***, $P < 0.001$). or $IRBIT^{KO}$ cells (*, $P < 0.05$). Acute application of 2-APB (100 µM) significantly reduced SOCE in all cells (####, $P < 0.0001$). Two-way ANOVA with Tukey's multiple comparisons test. Each bar represents the mean (± SD) of four separate experiments done in triplicate.

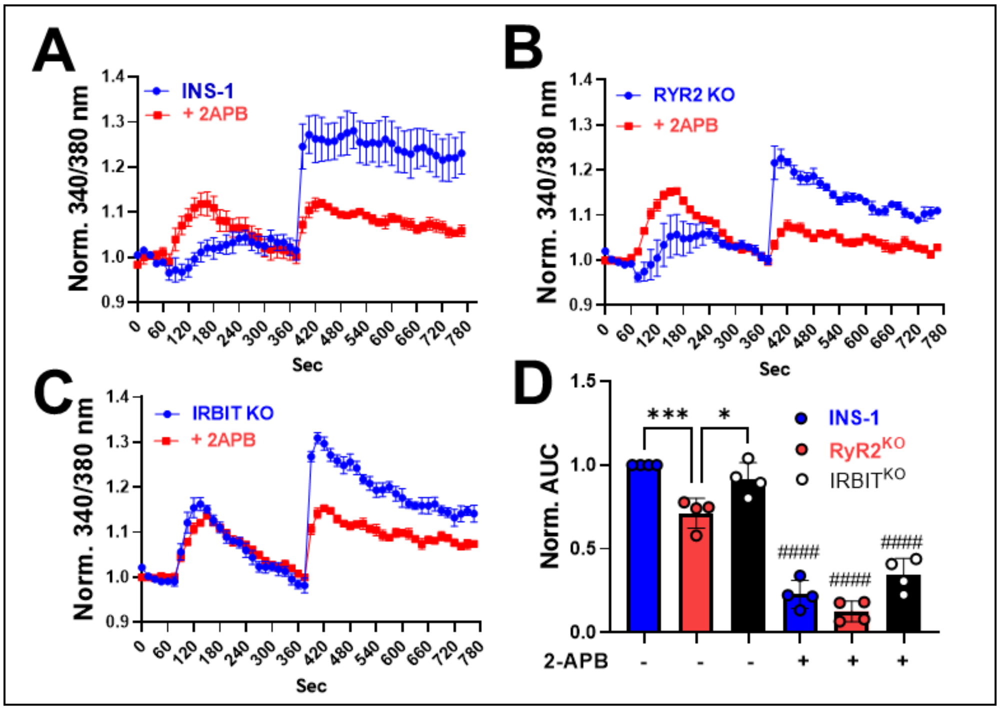

**Figure S2. Deletion of RyR2 or IRBIT increases peak current density independently of extracellular perfusion or extracellular solution composition in INS-1 cells. A)** Peak current density of INS-1, $RyR2^{KO}$, and $IRBIT^{KO}$ cells measured using the solution set given in Materials and methods ($Na^+/Ba^{2+}$) with or without perfusion with extracellular solution using a Biologic RSC 160 solution exchanger (***, $P < 0.001$; ****, $P < 0.0001$; One-way ANOVA with Tukey's multiple comparisons test). NOTE: data for $Na^+/Ba^{2+}$+ perfusion is the same as presented in Figure 2C. $Na^+/Ba^{2+}$ (no perfusion): INS-1 n = 14; $RyR2^{KO}$ n = 22; $IRBIT^{KO}$ n = 20. **B)** Peak current density of INS-1 and $RyR2^{KO}$ cells measured a Tris-$Ba^{2+}$ solution set with perfusion with extracellular solution or in a Tris-$Ca^{2+}$ solution set with or without perfusion with extracellular solution. (****, $P < 0.0001$; un-paired Welch's t-test). Tris-$Ba^{2+}$ + Perfusion: INS-1 n = 15; $RyR2^{KO}$ n = 14. Tris-$Ca^{2+}$ (no perfusion): INS-1 n = 13; $RyR2^{KO}$ n = 16. Tris-$Ca^{2+}$ + Perfusion: INS-1 n = 12; $RyR2^{KO}$ n= 16. Data are shown as mean ± SE. Tris-$Ba^{2+}$ solutions: Extracellular: 150 mM Tris, 10 mM $BaCl_2$, 4 mM $MgCl_2$ adjusted to pH 7.3 with methanesulfonic acid; Intracellular: 130 mM N-methyl-D-glucamine, 60 mM HEPES, 10 mM ethylene glycol-bis (β-aminoethyl ether)-N,N,N',N'-tetraacetic acid, 2 mM ATP, 1 mM $MgCl_2$, pH adjusted to 7.3 with methanesulfonic acid. Tris-$Ca^{2+}$ solutions were the same as Tris-$Ba^{2+}$ except that 10 mM $CaCl_2$ was substituted for 10 mM $BaCl_2$ in the intracellular solution.

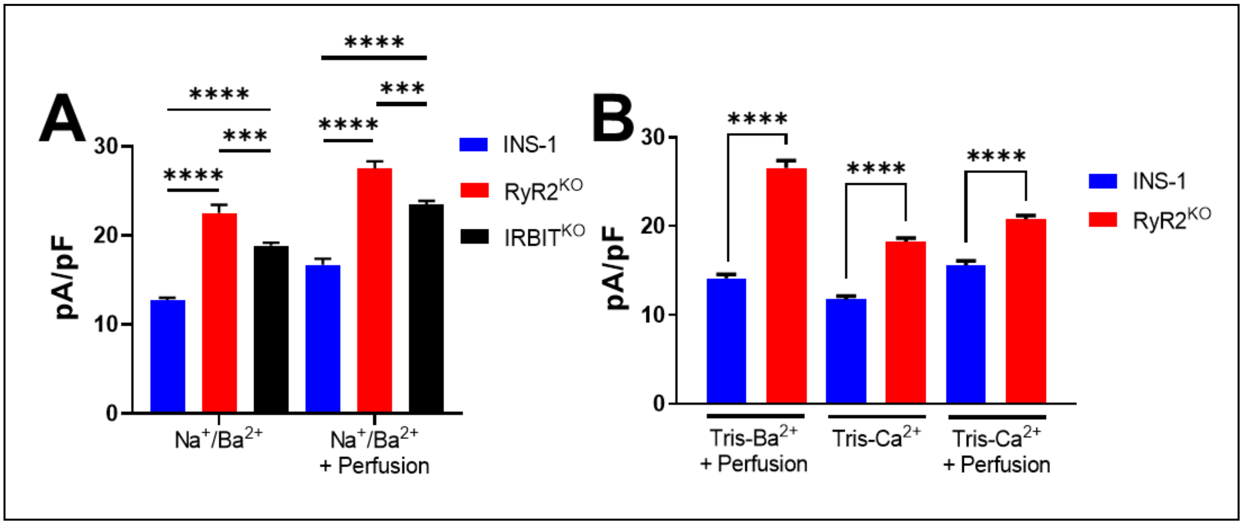
